# Unsupervised machine learning reveals key immune cell subsets in COVID-19, rhinovirus infection, and cancer therapy

**DOI:** 10.1101/2020.07.31.190454

**Authors:** Sierra M. Barone, Alberta G.A. Paul, Lyndsey M. Muehling, Joanne A. Lannigan, William W. Kwok, Ronald B. Turner, Judith A. Woodfolk, Jonathan M. Irish

**Affiliations:** Department of Cell and Developmental Biology, Vanderbilt University, Nashville, TN, USA; Vanderbilt-Ingram Cancer Center, Vanderbilt University Medical Center, Nashville, TN, USA; Allergy Division, Department of Medicine, University of Virginia School of Medicine, Charlottesville, VA, USA; Department of Microbiology, Immunology, and Cancer Biology, University of Virginia School of Medicine, Charlottesville, VA, USA; Benaroya Research Institute at Virginia Mason, Seattle, WA, USA; Department of Pediatrics, University of Virginia School of Medicine, Charlottesville, VA, USA; Department of Pathology, Microbiology and Immunology, Vanderbilt University Medical Center, Nashville, TN, USA

**Author notes:** Denotes equal contribution (Sierra M. Barone, Alberta G.A. Paul, Lyndsey M. Muehling). Corresponding authors (J.M.I.); (J.A.W.).

**Keywords:** Machine learning, rhinovirus, COVID-19, cancer immunology, systems immunology, human immune monitoring, single cell, spectral flow cytometry, mass cytometry

## Abstract

For an emerging disease like COVID-19, systems immunology tools may quickly identify and quantitatively characterize cells associated with disease progression or clinical response. With repeated sampling, immune monitoring creates a real-time portrait of the cells reacting to a novel virus before disease specific knowledge and tools are established. However, single cell analysis tools can struggle to reveal rare cells that are under 0.1% of the population. Here, the machine learning workflow Tracking Responders Expanding (T-REX) was created to identify changes in both very rare and common cells in diverse human immune monitoring settings. T-REX identified cells that were highly similar in phenotype and localized to hotspots of significant change during rhinovirus and SARS-CoV-2 infections. Specialized reagents used to detect the rhinovirus-specific CD4^+^ cells, MHCII tetramers, were not used during unsupervised analysis and instead ‘left out’ to serve as a test of whether T-REX identified biologically significant cells. In the rhinovirus challenge study, T-REX identified virus-specific CD4^+^ T cells based on these cells being a distinct phenotype that expanded by ≥95% following infection. T-REX successfully identified hotspots containing virus-specific T cells using pairs of samples comparing Day 7 of infection to samples taken either prior to infection (Day 0) or after clearing the infection (Day 28). Mapping pairwise comparisons in samples according to both the direction and degree of change provided a framework to compare systems level immune changes during infectious disease or therapy response. This revealed that the magnitude and direction of systemic immune change in some COVID-19 patients was comparable to that of blast crisis acute myeloid leukemia patients undergoing induction chemotherapy and characterized the identity of the immune cells that changed the most. Other COVID-19 patients instead matched an immune trajectory like that of individuals with rhinovirus infection or melanoma patients receiving checkpoint inhibitor therapy. T-REX analysis of paired blood samples provides an approach to rapidly identify and characterize mechanistically significant cells and to place emerging diseases into a systems immunology context.

## Introduction

Systems immunology offers a new way to compare how an individual patient’s cells respond to treatment or changes during infection ^1, 2^. However, systems immunology and computational analysis tools were primarily designed to track major cell populations representing >1% of a sample. Viral immune and cancer immunotherapy responses can include mechanistically important and extremely rare T cells that proliferate rapidly over the course of days but as an aggregate exist as <0.1% of blood CD3^+^ T cells at their peak. These cells can be tracked genetically through clonal expansion, but may be lost in computational analyses focused on describing the global landscape of phenotypes. The specific expansion or contraction of phenotypically distinct cells may be a hallmark feature of key immune effectors and could reveal these cells without the need for prior knowledge of their identity or specialized tracking reagents like MHC tetramers.

The datasets tested here were all suspension flow cytometry, a data type where it is typical to have multiple snapshot samples of cells over time; however, an ongoing challenge in the field is to match or register cells to their phenotypic cognates between samples ^3–5^. Analysis algorithms typically rely on aggregate statistics for clustered groups of cells, but the process of grouping the cells works best with larger, established populations ^6–8^ and can depend on pre-filtering of cells by human experts ^1, 9^. Cytometry tools like SPADE ^10, 11^, FlowSOM ^12^, Phenograph ^13^, Citrus ^14^, and RAPID ^15^ generally work best to characterize cell subsets representing >1% of the sample and are less capable of capturing extremely rare cells or subsets distinguished by only a fraction of measured features. Tools like t-SNE ^16, 17^, opt-SNE ^18^, and UMAP ^19, 20^ embed cells or learn a manifold and represent these transformations as algorithmically-generated axes. For a biologist, these tools provide a way to organize cells according to phenotypic relationships that span multiple measured features, such as the proteins quantified on each of millions of cells in the datasets here. In addition to assisting with data visualization, these tools frequently reveal unexpected cells and facilitate their identification through manual or automated clustering ^6, 15, 16, 21–23^. Sconify ^24^ is one such tool that applies *k*-nearest neighbors (KNN) to calculate aggregate statistics for the immediate phenotypic neighborhood around a given cell on a t-SNE plot representing data from multiple cytometry samples. This approach to creating a population around every cell was a key inspiration for the Tracking Responders Expanding (T-REX) tool presented here, which applies KNN to every cell to pinpoint rare cells in phenotypic regions of significant change. In addition to combining UMAP, KNN, and MEM in a rapid, unsupervised analysis workflow for paired samples from one individual, T-REX contrasts with prior approaches in its specific focus on the regions of great difference between samples. This T-REX design is based on the observation that, in the absence of a perturbation such as disease or infection, adults tend to have a stable signature of blood cell abundances over weeks to months ^9, 25, 26^, and the hypothesis that short-term, dramatic changes in rare immune cell subsets will identify cells associated with exposure to an immunogenic agent, such as a virus.

Data types used to challenge the T-REX algorithm here included a new spectral flow cytometry study (Dataset 1) and three existing mass cytometry datasets (Dataset 2, Dataset 3, and Dataset 4). Mass cytometry is an established technique for human immune monitoring where commercial reagents presently allow 44 antibodies to be measured simultaneously per cell ^1, 27, 28^. Spectral flow cytometry is gaining attention in human immune monitoring as it generates data that compares well to mass cytometry ^27, 29^. Spectral flow cytometers collect cells at around 10-fold the number of cells per second as mass cytometers. While the availability of spectrally distinct antibody-fluorochrome conjugates imposes some practical limits on spectral flow cytometry at present, established panels like the one in Dataset 1 measure ~30 features per cell with excellent resolution, and that capacity is expected to roughly double in the next few years as recent work has demonstrated 40 features ^30^. Spectral flow cytometry is thus well-matched to studies of very low frequency cells, as was the case in Dataset 1, where a goal was to computationally pinpoint hundreds of virus-specific T cells in datasets of over 5 million collected cells.

Datasets 1 and 2 were from individuals infected with two different respiratory viruses, rhinovirus or SARS-Cov-2, respectively. Respiratory viruses are ubiquitous and while some, like rhinovirus, are generally benign, they nonetheless pose risks to patients with underlying chronic health conditions. The common colds associated with rhinovirus are characterized by shifts in very rare virus-specific cells in the blood ^31, 32^. In contrast, novel respiratory viruses, such as SARS-CoV-2, the coronavirus causing COVID-19, continue to emerge that enact high morbidity and mortality, even among healthy subjects. Understanding the immune response to such viruses is vital to treatment and vaccine design, and there has been rapid progress applying human immune monitoring to COVID-19 patients ^25, 33^. An ongoing challenge in the field is to quantitatively compare novel diseases, like COVID-19, to other disease states and immune responses. T cells are pivotal to such responses. Severe COVID-19 has been linked to a pathogenic “cytokine storm” in which cellular immune responses likely play a crucial role ^34^. Nonetheless, in the case of both rhinovirus and COVID-19, it is clear that host factors are a key determinant of the degree of the T cell response ^25, 31^. Datasets 3 and 4 were from cancer patients that included melanoma patients being treated with α-PD-1 checkpoint inhibitor therapies or acute myeloid leukemia patients undergoing induction chemotherapy. By tracking the CD4^+^ T cells that expand rapidly during infections and respond to immunotherapy, it may be possible to pinpoint or therapeutically guide cells into helpful vs. harmful roles or niches. Overall, a goal of this study was to develop an automated, quantitative toolkit for immune monitoring that would span a wide range of possible immune changes, identify and phenotype statistically significant cell subsets, and provide an overall vector of change indicating both the direction and magnitude of shifts, either in the immune system as a whole or in a key cell subpopulation.

## Results

We report here Tracking Responders Expanding (T-REX), a novel unsupervised machine learning algorithm for characterizing cells in phenotypic regions of significant change in a pair of samples (Figure 1). The primary use case for developing the T-REX algorithm was a new dataset from individuals infected with rhinovirus, where changes in the peripheral immune system are expected in very rare memory cells responding directly to the virus (Dataset 1). Infection with rhinovirus is known to induce expansion of circulating virus-specific CD4^+^ T cells in the blood, and a key feature of the new rhinovirus dataset here is that rare and mechanistically important virus-specific CD4^+^ T cells were marked with MHC II tetramers in the context of multiple other T-cell markers. The T-REX algorithm was blinded to tetramers during analysis so that they could subsequently be used to test algorithm performance. In addition, T-REX was tested with paired samples from patients with moderate or severe COVID-19 (Dataset 2), melanoma patients being treated with α-PD-1 checkpoint inhibitor therapies (Dataset 3), and acute myeloid leukemia patients undergoing induction chemotherapy (Dataset 4). These datasets were used to determine whether the T-REX algorithm functions effectively across a spectrum of human immune monitoring challenges and to see how the algorithm performs when changes are restricted to rare cell subsets, as in Dataset 1 and Dataset 3, or when many cells may be expanding or contracting, as in Dataset 2 and Dataset 4.

**Figure 1.**
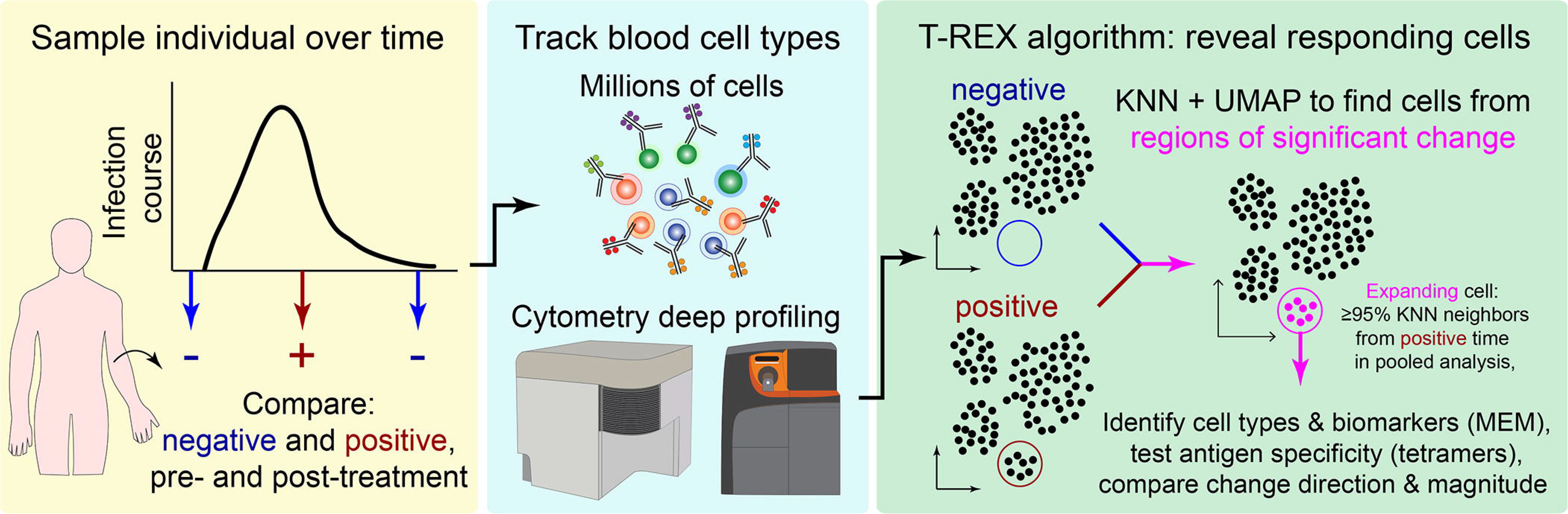
Tracking Responders EXpanding (T-REX) algorithm identifies rare cells based on significant expansion or contraction during infection or treatment. Graphic of the Tracking Responders Expanding (T-REX) workflow. Data from paired samples of blood from a subject are collected over the course of infection and analyzed by high dimensional, high cellularity cytometry approaches (e.g., Aurora or CyTOF instrument, as with datasets here). Cells from the sample pair are then equally subsampled for UMAP analysis. A KNN search is then performed within the UMAP manifold for every cell. For every cell, the percent change between the sample pairs is calculated for the cells within its KNN region. Regions of marked expansion or contraction during infection are then analyzed to identify cell types and key features using MEM. For some datasets, additional information not used in the analysis could be assessed to determine whether identified cells were virus-specific. Finally, the average direction and magnitude of change for cells in the sample was calculated as an overall summary of how the analyzed cells changed between samples.

### T-REX identifies cells in phenotypically distinct regions of significant change

For the rhinovirus challenge study in Dataset 1, sample pairs available for T-REX included cells taken immediately prior to intranasal inoculation with virus (i.e., pre-infection, day 0), as well as those during (day 7) or following inoculation (day 28). Cells were subsampled equally from each timepoint and then concatenated for a single UMAP specific to each analysis pair. UMAP axes were labeled to indicate they were specific to a comparison for a given individual (Figure 2). Thus, each UMAP comparison was a new run of the algorithm. Although it is also possible to map all sample times or all individuals into a single UMAP for analysis, a key goal here was to imagine a minimal T-REX use case with only a pair of samples from one individual. The features selected for UMAP analysis were intentionally limited to surface proteins in order to test whether suitable features for live cell fluorescence activated cell sorting (FACS) could be identified. Following UMAP, each cell was used as the seed for a KNN search of the local neighborhood within the UMAP axes (i.e., the KNN search was within the learned manifold, as with the analysis in Sconify ^24^ or RAPID ^15^). The k-value for KNN was set to 60 as a starting point based on prior studies and later optimized. For each cell, the KNN region could include cells from either time chosen for analysis, and the percentage of each was calculated to determine the representation of each sampled time in a cellular neighborhood. When cells in regions of expansion (≥95% of cells in the KNN region from one sampling time) were clustered together in one phenotypic region of the UMAP, they were considered a ‘hotspot’ of significant change. Cells in change hotspots were aggregated and the phenotype automatically characterized using Marker Enrichment Modeling (MEM) ^21^. MEM labels here indicated features that were enriched relative to a statistical null control on a scale from 0 (no expression or enrichment) to +10 (greatest enrichment). Ultimately, T-REX and MEM were used to reveal hotspots of ≥95% change and assign a label that could be used by experts to infer cell identity.

**Figure 2.**
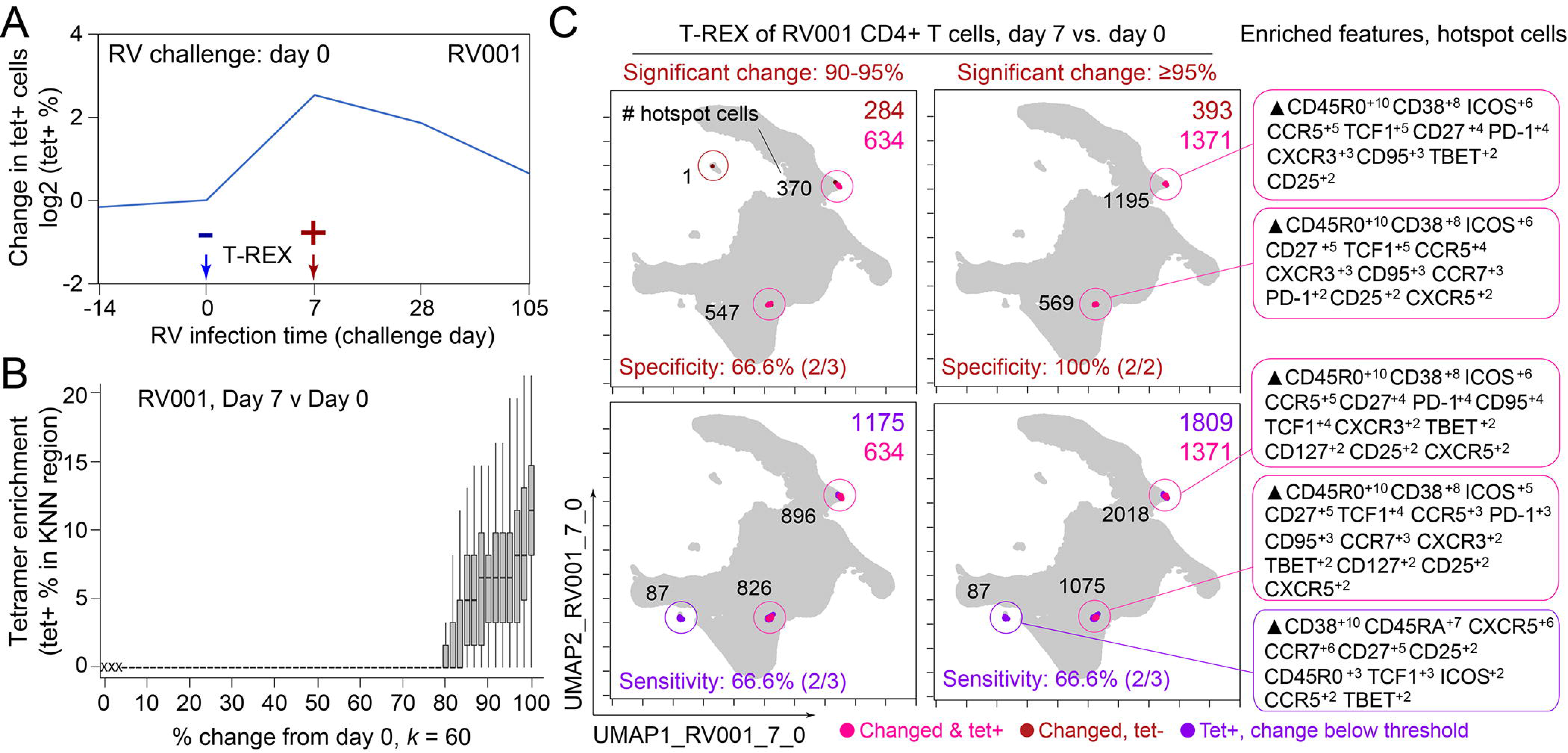
T-REX identifies molecular signatures of CD4+ T cells that are expanded during acute rhinovirus infection and enriched for virus-specific cells. A subject (RV001) was experimentally infected with rhinovirus (RV-A16) and CD4+ T cell signatures monitored by spectral flow cytometry in conjunction with tetramer staining during the course of infection. (A) Fold change in the number of tetramer-positive cells (log2) after rhinovirus challenge on day 0. (B) Data showing the percentage of tetramer+ cells in each cell’s KNN region (where k = 60) plotted against the percentage change in its KNN region on day 7 vs. day 0. A statistical threshold of 80% or higher for the percentage change in KNN region corresponded to marked enrichment of tetramer+ cells at day 7. (C) UMAP plots with T-REX analysis of CD4+ T cells for day 7 vs. day 0 based on statistical thresholds of 90-95% change (left column) and ≥95% change (right column) in cell phenotypes. Pink and red colors denote regions of phenotypic change identified by T-REX. Numbers of tetramer+ cells within the cell’s KNN region captured in these areas of phenotypic change are denoted. Cells containing >5% tetramer+ virus-specific cells in the corresponding KNN region were labeled pink. Red cells denote a KNN region that was not enriched for tetramer+ cells, and purple cells denote a tetramer enriched region not captured by T-REX. Values in black indicate the actual number of tetramer+ cells in each circled hotspot of phenotypic change. MEM labels on the right indicate cell phenotypes of each hotspot.

In the human rhinovirus challenge study yielding Dataset 1, MHC class II tetramers were used to identify rhinovirus-specific CD4^+^ T cells with the goal of tracking phenotypic changes over the course of infection. Increases in tetramer^+^ cells on day 7 (Figure 2A) corresponded to the acute infection phase ^31^. This tetramer tracking system for virus-specific T cells provided an opportunity to test whether the cells identified by T-REX were biologically significant by leaving the tetramer stain features out of the computational analysis (i.e., not using tetramers to make the UMAP or in other parts of T-REX) and then testing to see whether hotspots of cellular change identified by T-REX were statistically enriched for tetramer^+^, virus-specific cells. In the example subject shown, the pairwise comparisons used in T-REX analysis included CD4^+^ T cells from day 0, immediately prior to rhinovirus infection, and day 7, a well-studied time point at which rare, virus-specific CD4^+^ T cells are observed at higher frequencies ^31^. This trajectory of virus-specific cell expansion was confirmed by a peak in the log_2_ fold change in the frequency of tetramer^+^ CD4^+^ T cells (Figure 2A). Applying T-REX to the rhinovirus data revealed that KNN regions with expansion from day 0 to day 7 were greatly enriched for tetramer^+^ cells, as compared to regions with less expansion (Figure 2B). UMAP axes were labeled as UMAP_RV001_7_0 to denote this UMAP analyzed day 0 and day 7 for individual RV001 (Figure 2C). Regions of contraction were observed but were not enriched for tetramer^+^ cells, except in the case of one individual, RV007, studied here (Figure 3). Notably, two of the eight study subjects challenged with rhinovirus were not infected (RV002 and RV003); all other individuals were infected (Supplemental Table 1).

**Figure 3.**
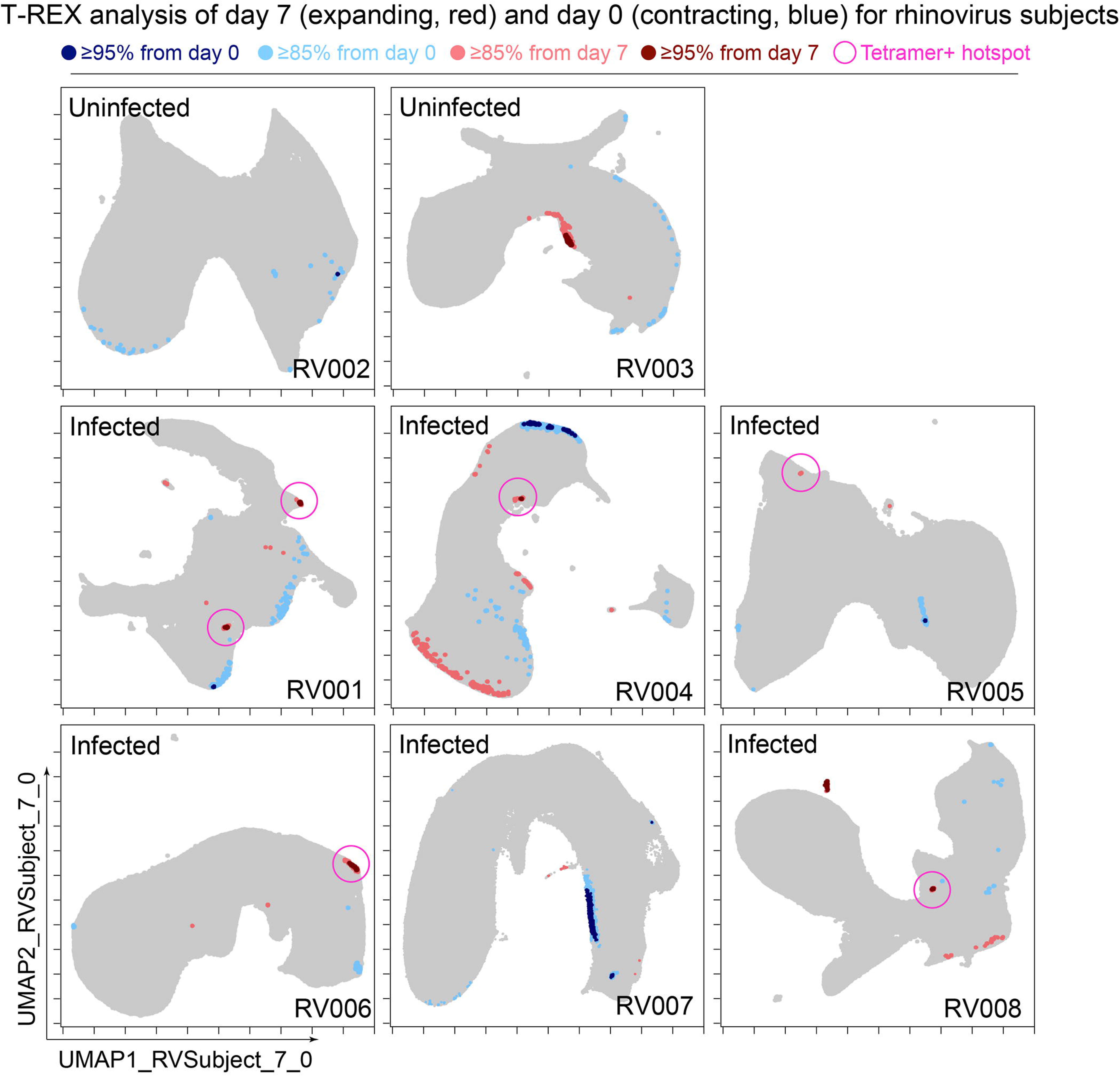
Cells in regions of significant change between day 0 and day 7 were typically in tetramer^+^ hotspots. T-REX plots of regions of significant change (blue and red) are shown on UMAP axes for CD4+ T cells from 8 rhinovirus challenge study individuals. Solid pink circles indicate tetramer+ hotspots that also contained cells that were in regions of marked expansion ≥85%

A key question for the T-REX algorithm is where to set a statistical cutoff for what is considered to be a biologically significant amount of expansion. Two change cutoffs were tested with subject RV001, ≥90% and ≥95% (Figure 2C). Using a cutoff of ≥95% identified 2/2 (100%) tetramer+ hotspots of change for RV001 and did not identify any additional regions that were not tetramer hotspots, whereas the ≥90% cutoff identified both tetramer+ hotspots and an additional tetramer-hotspot (Figure 2C, *top*). Thus, ≥95% represented a stringent cutoff that still captured biologically significant cells. An analysis of tetramer enrichment as a function of percentile of expansion from day 0 to day 7 (Figure 2B) showed that tetramer^+^ cells were not commonly observed to be in local neighborhoods around cells with change below 80% in their KNN region. In contrast, above 90% change, the median CD4^+^ T cell had 10% or more tetramer^+^ neighbors around it in the KNN region (Figure 2B). Thus, only regions of 80% or more expansion from day 0 to day 7 were enriched for tetramer^+^ CD4^+^ T cells in study individual RV001.

### A *k*-value of 60 effectively identified immune hotspots in T-REX

A critical question for KNN analysis is the value of k, the number of neighbors to assess. While it is useful to have a lower *k*-value as the analysis will complete more quickly, increasing the *k*-value might better represent the phenotypic neighborhood or be more statistically robust. To assess how *k*-value impacted detection of cells in regions of change and the degree to which these cells were virus-specific in rhinovirus challenge Dataset 1, the k-value was systematically changed. In example case RV001, an optimal *k* was determined to be an inflection point in a graph of the average tetramer enrichment (y-axis, Figure 4) versus increasing values of *k* (x-axis, Figure 4). To calculate this curve, a KNN search was repeated while increasing *k* in steps from 0 to 300 for every cell in each sampling. This analysis was performed for all tetramer^+^ cells from day 7 (dark purple, Figure 4), all tetramer^+^ cells from day 0 (light purple, Figure 4), and, as a negative control, random tetramer negative cells from day 7 (black, Figure 4). Within each of these neighborhoods, tetramer enrichment was calculated. This approach identified the inflection point of the tetramer^+^ density curve as *k* = 70 for RV001 (Figure 4). In further analysis of the remaining infected rhinovirus subjects, optimal k values ranged from 40 to 80. A k value of 60 was chosen and used in all other analyses of rhinovirus subjects (Figure 3), as well as Datasets 2, 3, and 4 described below.

**Figure 4.**
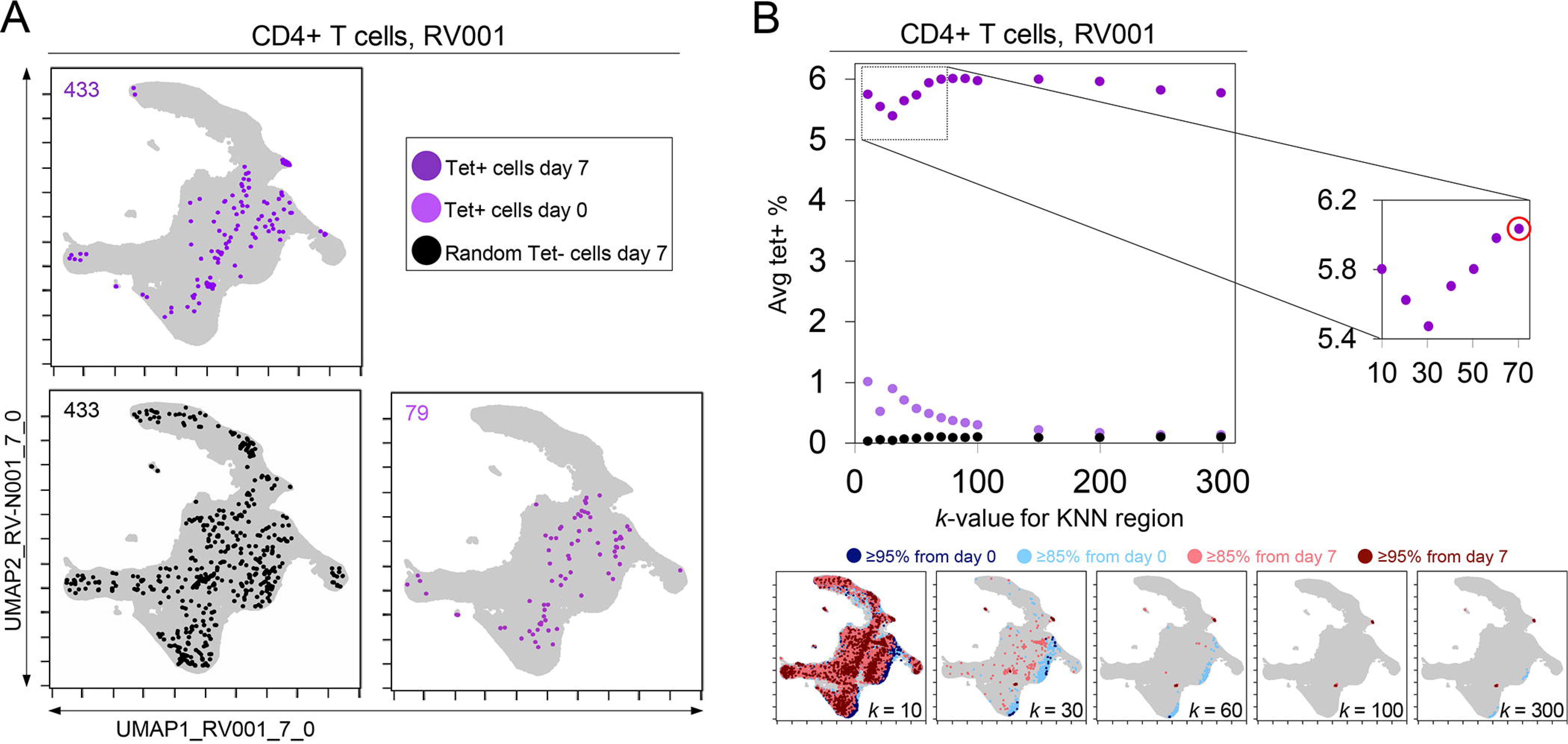
KNN analysis around tetramer^+^ cells reveals an optimized *k*-value at the inflection point of the tetramer density curve. (A) Tetramer^+^ cells from day 7 (dark purple) or from day 0 (light purple) and random tetramer^-^ cells from day 7 (black) are shown overlaid on a common UMAP plot. The number of cells for each group is shown in the upper left of each plot. (B) Average tetramer enrichment is shown for increasing *k*-values in repeated KNN analysis of the cells. The inflection point of the resulting curve is circled in red at *k* = 70, which was the optimized *k*-value for KNN implemented as in T-REX for subject RV001. The T-REX plots on the UMAP axes are shown for various *k*-values.

### Regions of significant change contained rhinovirus-specific CD4+ T cells in Dataset 1

The association between regions of change and enrichment for virus-specific cells observed in the example subject shown (Figure 2B) was observed in five infected rhinovirus subjects; tetramer^+^ CD4^+^ T cells were not enriched in KNN regions around cells that had not expanded from day 0 to day 7 (1 infected, 2 uninfected; Supplemental Figure 1). This observation suggested that cutoffs at the 5^th^ and 95^th^ percentile would accurately capture cells representing phenotypic regions with significant change over time. In addition, 15^th^ and 85^th^ percentiles were chosen as cutoffs to capture a more moderate degree of change and track cells that might still be of interest but not from regions experiencing significant change. The remaining cells in phenotypic regions between the 15^th^ and 85^th^ percentiles were not considered to have not changed significantly in the context of these studies.

Going forward, it was of interest to determine how often regions of significant change (i.e., the 95^th^ and 5^th^ percentile cutoffs) would contain tetramer^+^ CD4^+^ T cells in different individuals participating in the rhinovirus challenge study. Cells in regions of significant expansion (≥95%) were also from regions that were enriched for virus-specific cells in nearly all rhinovirus-infected individuals (4/6 at 95% cutoff, 5/6 at 85% cutoff) (Figure 2, Supplemental Figure 1, Figure 3). Thus, by focusing specifically on cells in regions representing the most change over time, T-REX analysis revealed subpopulations containing virus-specific cells. This highlights the ability of T-REX to pinpoint such cells without the use of antigen-specific reagents. Following T-REX, MEM analysis was performed using all available features, including intracellular features not used to define the UMAP space (TCF1, TBET, and Ki-67). The phenotype of the regions of significant change enriched for virus-specific cells was quantitatively described with MEM scores (hotspot 1: ▲CD45R0^+10^ CD38^+8^ ICOS^+6^ CCR5^+5^ TCF1^+5^ CD27^+4^ PD-1^+4^ CXCR3^+3^ CD95^+3^ TBET^+2^ CD25^+2^; hotspot 2: ▲CD45R0^+10^ CD38^+8^ ICOS^+6^ CD27^+5^ TCF1^+5^ CCR5^+4^ CXCR3^+3^ CD95^+3^ CCR7^+3^ PD-1^+2^ CD25^+2^ CXCR5^+2^). The change hotspots thus contained activated memory cells (CD45RO+CD38+) that were notable for their early memory/stem-like T cell signature (TCF1+CD27+), as well as their expression of CCR5 and CXCR3, both of which are chemokine receptors found on rhinovirus-specific CD4+ T cells that respond during infection ^31, 32^.

To determine the sensitivity of this method, all tetramer+ regions were next reviewed, including those that did not meet the criteria for hotspots of significant change (Figure 2C, *bottom*). In analysis of RV001, 66.6% (2/3) of tetramer+ regions were captured, meaning there was one region with lower change that contained tetramer+ cells. However, there were only 87 cells in these missed regions compared to 896 cells and 826 cells in the regions with ≥95% expansion, confirming that T-REX captured the majority of virus-specific cells in the data set.

In the case of an emerging infectious disease, it may not be possible to have a pre-infection sample and it would be useful to know whether T-REX analysis of change between a peak of infection and a later time might also reveal virus-specific T cells. To test this idea, pairwise comparisons were performed with cells from day 7 following rhinovirus inoculation and at day 28 after inoculation (Figure 5). Strikingly, cells in phenotypic regions of significant change again were enriched for virus-specific, tetramer^+^ CD4^+^ T cells. The MEM values for these cells further identified them as CD45R0^+^ memory cells enriched for CD38, ICOS, CD27, TCF1, CXCR5, PD-1, and CD95 expression, a phenotype matching that of the cells identified in the day 0 to day 7 analysis for this individual (RV001, Figure 5).

**Figure 5.**
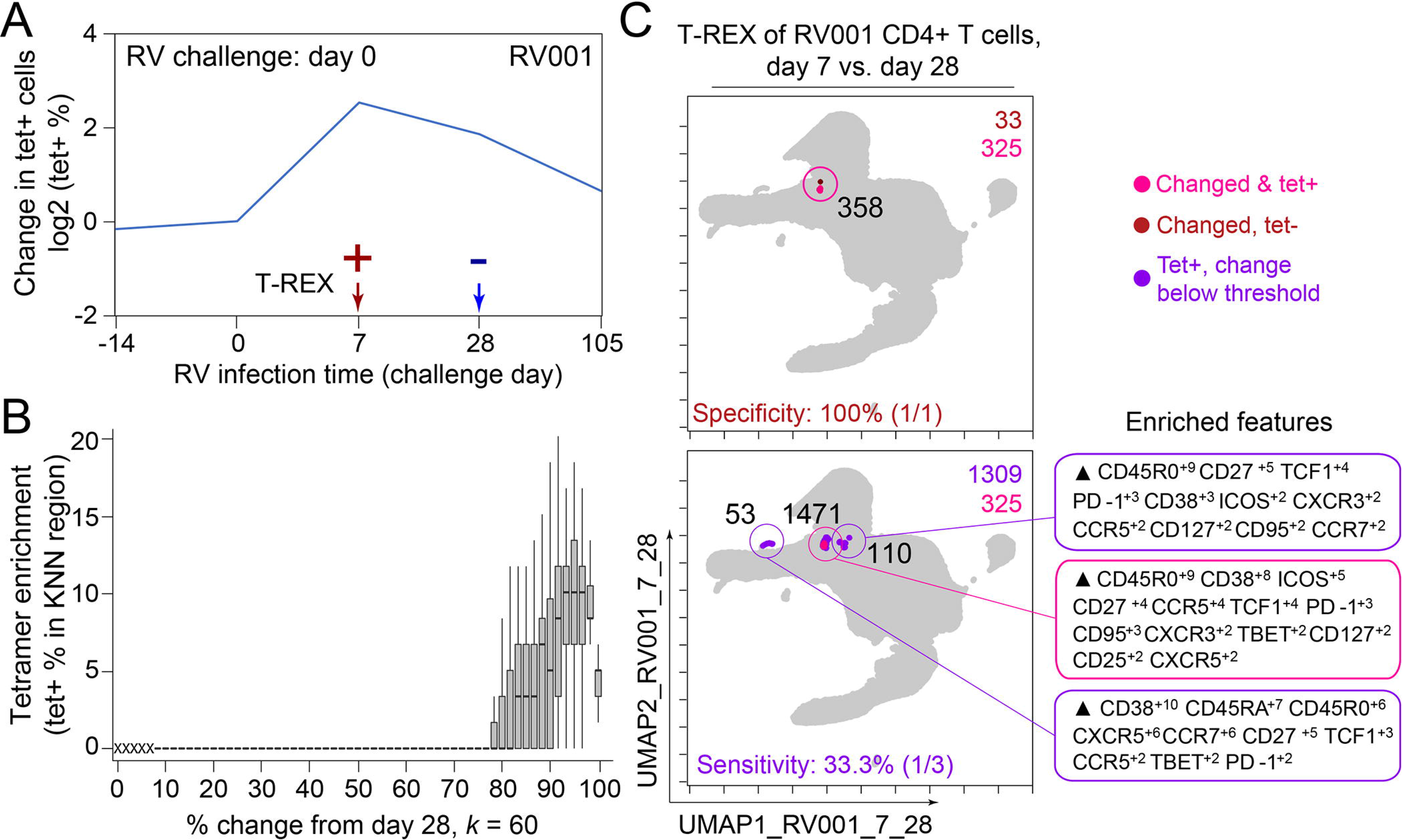
Infected cell phenotypes can be compared to cells taken after infection to reveal regions of expansion. (A) Fold change in the number of tetramer-positive cells (log2) after rhinovirus challenge on day 0. (B) Box and whisker plot show KNN regions in terms of expansion during infection represented by percent change as well as percent of tetramer positive cells for post-infection (day 28) and during infection (day 7). (C) UMAP plots for 95 percent change and 5 percent tetramer cutoffs. Cell count is in black as well as in the upper right of each UMAP plot. MEM labels are given for highly expanded and tetramer-enriched regions.

### Traditional Biaxial Gating of Cells Identified by T-REX and MEM Enriches for RV-Specific T Cells

Once identified by machine learning approaches, it can be useful to define a gating scheme that might be used to test whether computationally-defined cell subsets can be found using traditional gates. In addition, FACS sorting for live T cells could use surface antigens and biaxial gates to physically separate such cells, as is typical for interrogation *in vitro*. To test this idea computationally, the features enriched in MEM labels for cells identified by T-REX (MEM label average and standard deviation: ▲CD45R0^9±2.5^ CD38^7±2.0^ ICOS^6±1.4^ CCR5^4±1.7^ PD-1^4±0.9^ CD95^4±0.7^ CD27^3±1.6^ CXCR3^2±0.5^) were used to define a new gating strategy that used a single positive cutoff gate for each feature of CD4+ T cells (Live, Dump-, CD3+, CD4+) in the order CD45R0, CD38, ICOS, CCR5, PD-1, CD95, CD27, and CXCR3 (Supplemental Figure 2). At each gating step, the percentage of RV-specific cells was determined.

It is known that precursor frequencies of rhinovirus-specific CD4+ T cells are very low, even during active infection (0.0004L0.04% of CD4+ T cells, Supplemental Figure 3). The ‘virtual sort’ successfully enriched for rhinovirus tetramer+ cells in all infected subjects (0.89L9.25% of CD4+ T cells; Supplemental Figure 3). This is notable, considering that the consensus MEM label was generated from regions of ≥95% change, some of which did not include tetramers. Furthermore, this strategy was able to enrich for tetramer+ cells in the one infected subject for which T-REX was unable to identify tetramer hotspots (RV007), and one in which the tetramer+ hotspot only met a ≥85% threshold of expansion (RV005), suggesting that T-REX-derived sorting strategies can be broadly applied across cohorts, including subjects whose response may not reach the threshold of identification by T-REX. A minimal panel of ten markers (Live, Dump–, CD3+, CD4+, CD45R0+, CD38+, ICOS+, CCR5+, PD-1+, CD95+) was sufficient to achieve maximal tetramer enrichment.

Interestingly, gating for CD45R0 alone L the first MEM-enriched feature L did not significantly enrich for virus-specific T cells. Furthermore, the T-REX-derived sorting strategy failed to enrich for rhinovirus-specific T cells in uninfected subject, nor did it enrich for CD4^+^ T cells stained with a control influenza tetramer, confirming the specificity of this method (Supplemental Figure 3). Thus, a computational ‘virtual sort’ for the cells suggests that FACS gates could be drawn using the results of T-REX and MEM analysis. This result further confirms that the populations identified computationally also exist as populations that can be defined in traditional ways.

### T-REX tracking of direction and degree of change contextualizes diverse immune responses

The next goal was to test T-REX with additional data types and to contextualize the results from rhinovirus infection (Dataset 1) with changes that might be observed in other immune responses, such as another respiratory infection (Dataset 2), cancer immunotherapy (Datasets 3), or cancer chemotherapy (Dataset 4). To accomplish this, metrics for degree of change as well as direction of change in each sample were devised (Figure 6A). Degree of change was calculated as the sum of the percent of cells in the 5^th^ and 95^th^ hotspots of change. Direction of change was calculated as the difference between the number of cells in the 95^th^ and 5^th^ hotspots of change divided by the sum of the number of cells in the 95^th^ and 5^th^ hotspots of change. This way of looking at the data provided a method for comparing changes in many disease types. Rhinovirus subjects had small changes in samples over time with a median of 0.019% and an interquartile range (IQR) of 0.0006% (on a magnitude of change scale from 0 to 100%). Rhinovirus also had large directionality across all subjects either up or down, with a median of 0.029 and an IQR of 2.00 on the directionality scale from −1.00 to +1.00. Thus, rhinovirus displayed an extremely low magnitude of change, as very rare cell subsets were responding, and the direction of this change was typically fairly high or low for a given individual (i.e., the changes were not balanced and tended to represent marked expansion or contraction in the rare subsets that changed; Figure 6).

**Figure 6.**
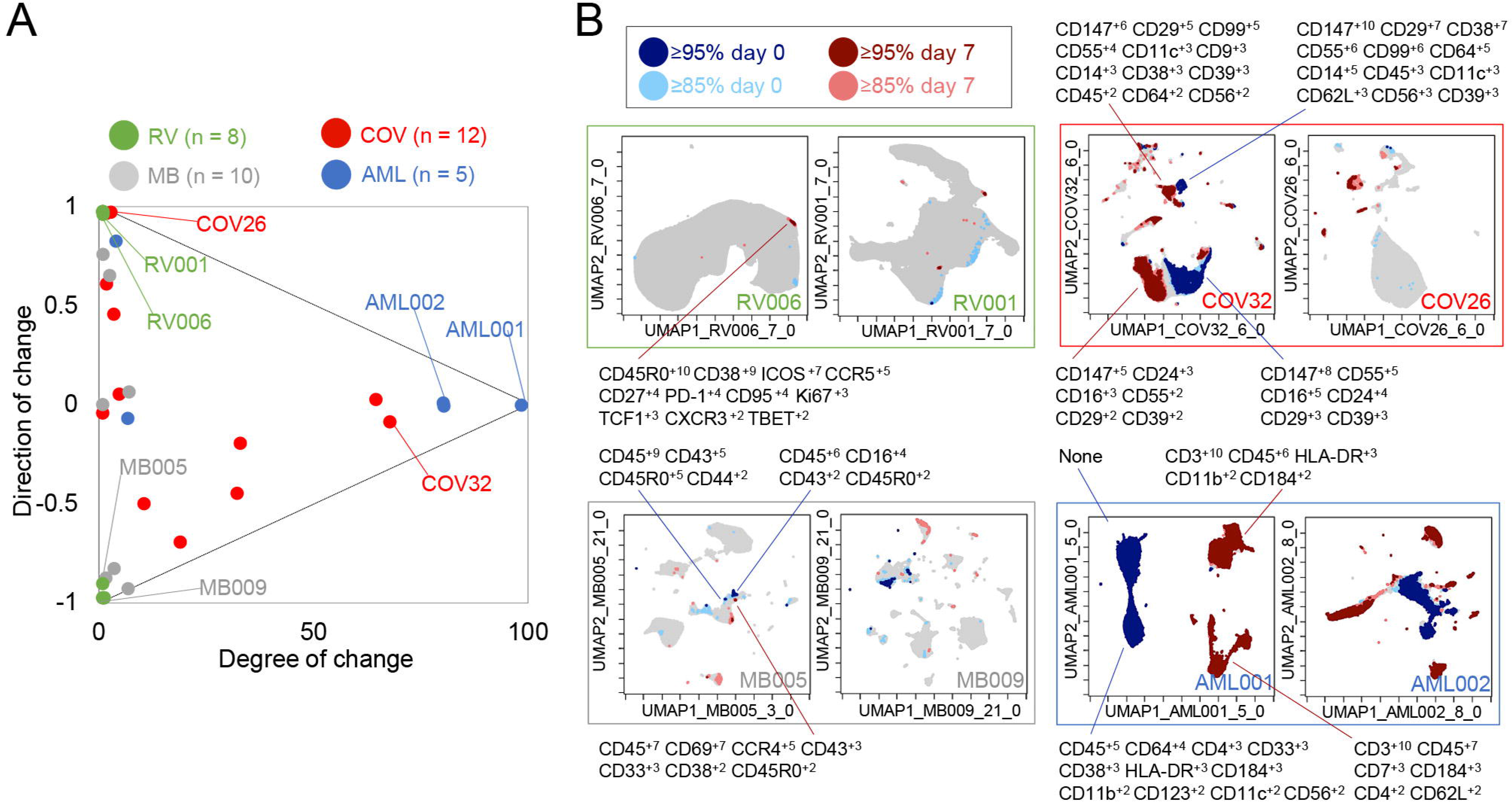
Mapping degree and direction of change for 5th & 95th hotspots reveals disease-specific patterns. (A) Degree of change and direction of change from T-REX analysis in a time point comparison shown for AML (day 5/8 vs day 0), COVID (day 1/3/4/5/6/7 vs day 0), MB (day 21/35 vs day 0), and RV (day 7 vs day 0) samples. (B) Example T-REX plots are shown for each disease type analyzed. Degree of change shown in red and blue with red showing regions of expansion over time compared to the blue representing regions of contraction over time. MEM label given for change hotspots in the left example in each sample type.

### Regions of change included cells expressing CD147 and CD38 in COVID-19 Dataset 2

Next, T-REX was applied to Dataset 2, a mass cytometry study of longitudinal collection of blood from patients with COVID-19 ^33^. This study originally contained data for 39 total patients, of which 12 patients had accessible mass cytometry data with at least two blood samples over time. For each patient, the day 0 timepoint and the closest sampled timepoint to day 7, were used for pairwise comparison using T-REX. The COVID-19 samples varied from <1% to 68% in terms of degree of change, with a median of 6.86% and an IQR of 30.4%. The directionality of change was near zero, with a median of −0.00880 and IQR of 0.773. Thus, the blood of COVID-19 patients could display marked changes or little change. Notably, the changes <5% were generally positive (median directionality of 0.55, N = 6), whereas the COVID-19 patient cell populations experiencing change >5% typically decreased between day 0 to day 7 (median directionality of −0.33, N = 6).

In T-REX analysis looking at change on the UMAP axes, patients with significant change were apparent due to large islands of cells being painted dark red or dark blue, indicating ≥95% change between paired days (Figure 6). These cell populations were clustered and separated into populations representing day 0 or the later time near day 7, and MEM labels calculated in order to assess the identity and phenotypic changes. For example, patient COV26 saw little change (magnitude of 2.02%) and this was almost entirely expansion (directionality of 0.99). The largest population experiencing significant change from COV26 decreased over time and had a MEM phenotype of CD147^+10^ CD99^+8^ CD29^+6^ CD38^+4^ CD55^+3^ CD14^+2^ CD39^+2^ CD64^+1^ CD56^+1^ CD8a^+1^, indicating it was a CD14^+^ myeloid cell subset with high expression of CD147/Basigin. The phenotypes for all automatically identified clusters of cells that expanded or contracted greatly, and the degree and direction of change for each COVID-19 patient from Dataset 2 is listed in Supplemental Table 2. These reference phenotypes should be comparable to those in other studies of COVID-19, and a meta-analysis of phenotypes could use quantitative analysis of MEM labels to compare these highly expanding and contracting cells.

T-REX was also applied to only CD4^+^ T cells from patients with sufficient T-cell counts (10 out of the 12 patients as described above). Of the 10 COVID-19 patients available for analysis, 5 individuals had at least one hotspot of great change, as revealed by T-REX, in CD4^+^ T cell specific analysis (Supplemental Figure 4, Supplemental Table 2). Analysis of the COVID-19 CD4+ T cell hotspot phenotypes using root mean square deviation (RMSD) analysis (Diggins et al., 2018; Diggins et al., 2017; Greenplate et al., 2019) identified three phenotypic groups. One of these groups was a set of closely related T cell subsets from one individual, patient COV32, and the aggregate MEM label for this population was (MEM label average and standard deviation: ▲CD57^+8±0.8^ CD99^+9±1.4^ CD29^+7±0.5^ CD147^+6±0.5^ CD43^+5±0.6^ CD45^+4±0.3^ CD3^+4±0.5^ CD81^+4±0.4^ CD52^+4±0.3^ CD49d^+3±0.5^ CD45RA^+3±2.3^ CD5^+3±1^ CD56^+2±1.5^, Supplemental Figure 3). Another phenotype of CD4+ T cells was consistently observed in those COVID-19 patients where T-REX revealed a hotspot. Of the 5 patients where T-REX identified a CD4+ T cell hotspot, 4 of the patients had a hotspot matching the aggregate phenotype (MEM label average and standard deviation: ▲CD147^+9±0.8^ CD99^+9±1.3^ CD29^+8±1.3^ CD45^+4±2^ CD3^+4±0.7^ CD38^+4±1.8^ CD49d^+3±1.6^ CD52^+3±1^ CD27^+3±2.1^ CD28^+3±0.8^ CD81^+2±1.3^ CD62L^+2±1.2^ CD56^+2±0.7^ CD5^+2±0.6^, Supplemental Figure 4). When comparing Day 0 to Day 6 (±3 days), this population of CD4^+^ T cells was observed to change significantly in patients COV24, COV29, COV32, and COV39 (4 of the 5 with a CD4^+^ T cell hotspot). The features of this subset could now be used to physically separate this population using FACS, highlighting a practical application. As a test of this idea, manual biaxial gating, as in standard FACS for physical separation of cell subset, was performed using the cell surface markers identified by MEM as most enriched, as in the prior analysis of cells from the rhinovirus study (Supplemental Figure 5). While CD4+ T cells only represented 5.5-14.0% of total cells, following MEM-based gating they were enriched to 49.6-83.3% of cells. Following expert gating, the MEM label of the resulting population was: ▲CD147^+9±0.6^ CD99^+8±1^ CD29^+7±0.6^ CD38^+7±0.8^ CD27^+5±0.8^ CD45^+4±1.2^ CD3^+4±0.6^ CD49d^+3±0.5^ CD81^+3±0.7^ CD52^+3±0.7^ CD28^+3±0.7^ CD62L^+2±0.8^ CD56^+2±0.5^ CD5^+2±0.6^, which closely matched the computationally identified cells.

### T-REX reveals immune cell changes during cancer therapies in Dataset 3 and Dataset 4

T-REX was next tested on two previously published cancer immune monitoring studies representing a wide range of immune system changes, from modest to extensive. Dataset 3 consisted of mass cytometry analysis of peripheral blood mononuclear cells (PBMC) from melanoma patients treated with anti-PD-1 ^9^. This well-studied dataset primarily includes melanoma patients whose blood had modest, subtle shifts in PBMC phenotypes over time. However, one patient in the set, patient MB-009, developed myelodysplastic syndrome (MDS) and experienced a great shift in blood immunophenotype in parallel with the emergence of a small population of blasts in PBMCs ^35^. Overall, when analyzed by T-REX, the melanoma samples in Dataset 3 for comparisons of day 21/35 versus day 0 had a small degree of change (median of 0.58% and an IQR 2.34%) with a varying directionality (median of −0.42 and an IQR of 1.46), confirming the subtle shifts in phenotypes as previously indicated. The great shift in peripheral immunophenotypes observed in MB-009 was confirmed with T-REX analysis when comparing the 6-week and 12-week times. Notably, at 6 weeks, the peripheral blast count was still below 5% ^35^, so T-REX detected a substantial change in subsets that were not driven solely by the emergence from the marrow of the MDS blasts.

Dataset 4 was chosen to represent large changes and included peripheral blood from AML patients treated with induction chemotherapy ^36^. The compared timepoints for the AML data in Dataset 4 were day 5/8 versus day 0. As expected, the majority of AML patients had a large degree of change in samples (median of 81.0% and an IQR of 75.2%) with little to no directionality to the change (median of −0.00250 and an IQR of 0.0173), meaning that there were massive changes in terms of both expansion and contraction over the course of treatment. MEM labels showed that the cells contracting in responder patients were the AML blasts, whereas the emerging cells were the non-malignant immune cells (Figure 6). AML samples with a degree of change >80% (AML001, AML002, AML004) came from patients with high blast count in the blood and complete response to treatment indicating the complete transformation of the immune environment after treatment. AML007, a patient with no blasts in the blood, had a degree of change of 5.97% over treatment. For AML003, a patient that did not respond to treatment, little change was seen from days 0 to 5 (degree of change = 3.19%) by means of T-REX analysis.

## Methods

### Generation of Dataset 1

Dataset 1 was a newly generated dataset of PBMCs obtained by longitudinal sampling of healthy volunteers who were challenged intranasally with RV-A16. The study was approved by the University of Virginia Human Investigations Committee, performed in accordance with the Declaration of Helsinki, and registered with ClinicalTrials.gov (NCT02796001). Informed consent was obtained from all study participants. Data were collected and processed at the University of Virginia. Sample collection times were defined by established kinetics of memory effector T helper cell responses ^31, 32^. Cells were stained with antibodies that target markers of naïve, memory and helper T cells (CCR6, ICOS, CXCR3, CD27, CCR5, TBET, CD45RA, CD45R0, CD95, CXCR5, TCF1, CCR7), and activation and proliferation (CD25, CD38, CD127, Ki-67, PD-1). The marker panel also included up to three MHCII/peptide tetramers to identify virus-specific CD4+ T cells ^37^. Data were collected using a 3-laser Aurora spectral flow cytometry instrument. Additional methodological details can be found on this article’s online supplement

### Data pre-processing

Before testing and evaluating the modular analysis workflow for rare cells, data preprocessing and QC of the data was done on all samples for all time points, which included spectral unmixing with autofluorescence subtraction, spill-over correction, and applying scales transformation. An arcsinh transformation was applied to the dataset with each channel having a tailored cofactor based on the instrument used to acquire the data as well as to stabilize variance near zero. Manual gating for clean-up of the data was done by an expert to exclude debris, doublets, and dead cells. As helper T cells were of interest for this RV study, the data analyzed was manually gated for CD3^+^ CD4^+^ T cells.

### T-REX algorithm

A modular data analysis workflow including UMAP, KNN, and MEM was developed in R and scripts for analysis of data in this manuscript are available online (https://github.com/cytolab/T-REX). The dimensionality tool used included UMAP, or Uniform Manifold Approximation and Projection. The default parameter settings for UMAP as found in the uwot package in R were used. Since UMAP analyses were specific to a given individual and pair of samples, UMAP axis were labeled to indicate the individual and comparison being made, as in ‘UMAP_RV001_07’, which indicated a comparison of day 0 and day 7 for individual RV001. The KNN search from the Fast Nearest Neighbors (FNN) package was used to find the nearest neighbors for a given cell. For this project, a KNN search was done for every cell using the low dimensional projection of the data as an input for the neighborhood search. The value for *k*, or the number of nearest neighbors, was determined by an optimization of tetramer enrichment within a neighborhood.

### MEM analysis of enriched features

Marker Enrichment Modeling from the MEM package (https://github.com/cytolab/mem) was used to characterize feature enrichment in KNN region around each cell. MEM normally requires a comparison of a population against a reference control, such as a common reference sample ^21^, all other cells ^15, 38^, or induced pluripotent stem cells ^9^. Here, a statistical reference point intended as a statistical null hypothesis was used as the MEM reference. For this statistical null MEM reference, the magnitude was zero and the IQR was the median IQR of all features chosen for the MEM analysis. Values were mapped from 0 enrichment to a maximum of +10 relative enrichment. The contribution of IQR was zeroed out for populations with a magnitude of 0.

### A Putative FACS Gating Strategy Based on T-REX Results

In order to assess the applicability of the T-REX algorithm in the development of follow-up FACS experiments, a sorting strategy was devised based on the results of T-REX and then tested computationally using Datasets 1 and 2 (Supplemental Figure 2, Supplemental Figure 5). To accomplish this, aggregate MEM scores of T-REX hotspots of ≥95% expansion were generated for each dataset. Cells were sequentially gated in the order of decreasing MEM feature enrichment, ending with a maximum set of 12 markers, reflecting common capabilities for cell sorting. In Dataset 1, the enrichment of tetramer+ cells was assessed in the populations resulting from putative sort gates, as compared with the total CD4+ T cell population. In Dataset 2, the enrichment of CD4+ T cells was assessed within the total cell population after similar gating using putative sort gates designed algorithmically based on the results of T-REX and MEM.

### Data availability and transparent analysis scripts

Datasets analyzed in this manuscript are available online, including at FlowRepository ^39^. COVID-19 Dataset 2 ^33^ (https://ki.app.box.com/s/sby0jesyu23a65cbgv51vpbzqjdmipr1), melanoma Dataset 3 ^9, 35^ (http://flowrepository.org/id/FR-FCM-ZYDG), and AML Dataset 4 ^9, 36^ (http://flowrepository.org/id/FR-FCM-ZZMC) were described and shared online in the associated manuscripts. Rhinovirus Dataset 1 is a newly generated dataset created at the University of Virginia available on FlowRepository (FR-FCM-Z2VX available at: http://flowrepository.org/id/RvFr2DwkDSym1BjCwd7ZJGNbl1ksyXT375C65F282JCjN30meLzqwn8G096D7H9D). Transparent analysis scripts for all four datasets and all presented results are publicly available on the CytoLab Github page for T-REX (https://github.com/cytolab/T-REX) and include open source code and commented Rmarkdown analysis walkthroughs.

## Discussion

A signature feature of the immune system is the ability of rare cells to respond to a stimulus by activating and proliferating, leading to rapid expansion of highly specialized cells that may share both a distinct phenotype and a clonal origin. Here, we report the design of a new algorithm that can identify and reliably phenotype biologically relevant cell types, including very rare cells, which respond in human disease. The T-REX algorithm was designed to capture phenotypic regions where significant change was occurring between a pair of samples from one individual. The fact that T-REX was able to identify the phenotype of cells whose regions were greatly enriched for virus-specific T cells in rhinovirus Dataset 1, highlighted its ability to pinpoint rare cells responding to infection, and closely matches what would be expected based on a current understanding of rhinovirus immunology ^31, 32^. Specifically, after rhinovirus challenge, expanded regions displayed molecular signatures consistent with activated memory (CD45RO+CD38+) and tissue trafficking (e.g., CCR5 enrichment in the MEM labels) that aligned with our previous findings for rhinovirus-specific CD4+ T cells using manual gating methods and a limited marker panel ^31, 40^. The algorithm also reliably identified memory phenotypes of cells responding to rhinovirus infection, thereby revealing its potential to track the evolution of memory responses *in vivo*, in addition to defining candidate signatures that might be probed in functional assays. Although comparative phenotyping across time was beyond the scope of this study, it will be of high interest in the future to determine whether the vector of change in specific subsets correlates with additional aspects of disease or complicating host factors, such as allergy and asthma.

The T-REX algorithm also revealed potential new research directions, as there were cells that one might predict would be virus-specific, based on the T-REX enrichment analysis, but which were not enriched for the specific tetramers available here (e.g., Figure 3, RV007). Genetic analysis for the clonal origin of cells in such regions might help to determine whether these cells correspond to a clonal response for which a tetramer was not available, or else another type of CD4^+^ T cell response that may or may not be related to rhinovirus infection, such as a “bystander” response. Additionally, it will be important to test whether this type of finding holds true for other well-studied viruses for which tetramers are available, such as influenza ^41^, and whether these findings extend to MHC class I tetramers and CD8 T cells. It was also striking that in the comparisons of day 7 to either day 0 (Figure 2 and Figure 3) or day 28 (Figure 5), only the expanding cells (red) were in regions that were also tetramer hotspots. However, despite the focus on expansion in the T-REX acronym, contracting cells will likely also be of biological significance in different disease settings (as with AML) or potentially at different time points during the course of an infection, for example as a result of egress from the circulation in the acute phase, or else transitions in memory and tissue-homing subsets that occur later. This aspect would also be expected to translate to different disease settings such as AML.

Extending the use of T-REX algorithm beyond rhinovirus further highlighted its ability to identify responding cells in a consistent manner across different subjects, and different disease settings. Indeed, it is notable that regardless of the disease context, the patient served as an effective baseline for comparison, and allowed T-REX to find phenotypically similar cells in individuals with different starting immune profiles. A central question in systems immune monitoring is to place newly emerging diseases into the context of other well-studied diseases and immune responses. In working to compare COVID-19 and rhinovirus, it became clear that a summary of change indicating both the direction and magnitude of shifts, was needed (Figure 6). This framework represents a way to summarize both broad populations of immune cells, like all CD45^+^ leukocytes, and key cell subpopulations, like CD4^+^ T cells. The striking changes observed in patients with moderate and severe COVID-19 were far beyond the subtle changes observed in individuals with rhinovirus and more closely matched the immune reprogramming observed in melanoma patients receiving checkpoint inhibitor therapy (Figure 6). A primary finding of T-REX analysis of Dataset 2 from the blood of COVID-19 patients was that some patients experienced very large changes in the blood, and that these changes were typically associated with more decreases than increases (Figure 6). This finding closely matches reported findings from others who observed a systematic reprogramming in many immune cell populations in severe COVID-19 patients ^25^. Also observed were T cell subsets with enrichment of CD38, PD-1, and CD95, as has also been reported. While disease severity is not available for individual patients from Dataset 2, it is known that all these cases were at least moderate or severe ^33^. It will be of interest to test the hypothesis that the more severe cases will be one of the two groups, either the patients with very little change and just expansion of cells, or those with more marked change and a general decrease of cells (Figure 6).

Notably, CD147/Basigin, was highly expressed on many cells that changed during infection and was observed to change greatly on some populations over time. CD147 has been proposed in pre-prints as both a binding partner for SARS-CoV-2 spike protein and a potential mechanism of cellular entry, although evidence is needed to support this controversial hypothesis ^42^. In the study of Dataset 2, the authors noted that immune responses were dominated by cells expressing CD38 and CD147 ^33^. In the T-REX analysis of the same Dataset 2, for the cells that were changing greatly, CD147 was sometimes present on cells from day 0 that decreased greatly and was lower or absent on cells that emerged only at later times (Figure 6). An example of this was seen in cells from patient COV40, for which the authors noted CD147 expression on effector subsets at 1 week and onwards. The cells pinpointed by T-REX as emerging at day 6 included B cells that expressed CD147 (e.g., CX3CR1^+8^ CD9^+8^ CD29^+8^ CD147^+5^ IgD^+3^ CD99^+3^ CD33^+1^ CD11c^+1^ HLA-DR^+1^ CD24^+1^, Supplemental Table 2), but the level of enrichment was lower than on myeloid cells from day 0 that decreased over time (e.g., CD147^+8^ CD29^+6^ CD55^+5^ CD38^+5^ CD99^+4^ CD64^+3^ CD62L^+2^ CD45^+1^ CD33^+1^ CD14^+1^, Supplemental Table 2). This pattern of decreased enrichment of CD147 on cells emerging after day 0 was seen on other patients (Supplemental Table 2) and is consistent with multiple explanations.

Overall, there was a strong downward trend in many of the markers and cell subsets in COVID-19 patients, suggesting either selection against cells expressing a high level of proteins, downregulation of expression of key surface markers like CD147, expansion of immature or abnormal cells, or extreme trafficking of cells into tissues. These potential outcomes cannot be distinguished from each other with the analysis here. The utility of the T-REX algorithm is primarily in generating these hypotheses automatically and in pinpointing cells with extreme behavior within the context of the patient as their own baseline. Given the large amounts of change (Figure 6) and the generally lower numbers of T cell subsets observed in COVID-19 than in healthy individuals (Supplemental Table 2), it may be the case that therapeutic stabilization of the immune system will be needed before virus-specific T cells will be identifiable with the T-REX method. It will be especially interesting to explore more mild cases of COVID-19 with this approach and determine whether the hotspots of change are truly virus-specific, analogous to the scenario with rhinovirus.

For the melanoma and AML cases presented here, the cohort sizes were not large enough to allow robust statistical comparison of patient response to degree or direction of change, although this information is available in the original studies ^9, 36^. Of the AML patients, those with a high magnitude of change (Figure 6) were also those that had a high blast count and were complete responders to induction therapy, suggesting that the change represents the overall “reset” of the immune system following chemotherapy. It will be of high interest to ask whether the identification of virus-specific T cells extends to populations of cells on checkpoint inhibitor therapy. The dynamics of regulatory cells may also be of interest, especially for autoimmunity, and it is possible, but not known, whether these cells will follow the same pattern as the CD4^+^ T cells in rhinovirus infection.

A major strength of the algorithm is that once cell regions of change are identified, the key features highlighted by T-REX and MEM can be used in lower parameter flow cytometry or imaging panels to provide further information, confirm findings, and physically isolate cells by FACS (Supplemental Figure 2). Thus, low parameter cytometry approaches may rely more on manual analysis methods and cell signatures that are determined *a priori*, and T-REX may provide a useful tool for narrowing in on such features using exploratory high dimensional data. The computational approach here emphasizes unsupervised UMAP and KNN clustering and uses statistical cutoffs to guide the analysis. Further optimization of the algorithm could include a stability testing analysis where the stochastic components of the algorithm are repeated to determine whether clusters or phenotypes are stable ^15, 43^. Overall, the unsupervised approach aims to diminish investigator bias and reveal novel or unexpected cell types. While unsupervised analysis tools have impacted high dimensional cytometry for at least a decade ^6, 7, 10, 16, 22^, T-REX is designed to capture both very rare and very common cell types and place them into a common context of immune change. The extremely rare T cells identified here are overlooked by other tools due to these tools typically needing clusters of cells representing at least 1% and generally more than 5% of the sample.

It is a central goal of systems immunology to map people with vastly contrasting immune system changes onto a common plot of change (as in Figure 6). The approach here goes beyond prior single measures of systematic change, such as Earth Mover’s Distance ^9, 44^, by including both direction and magnitude of change in one view of an individual’s immune response. This improvement proved useful for comparing settings with great change in many cell types (COVID-19 infection, AML chemotherapy responders) to settings with rare cells that specifically expanded or contracted (rhinovirus infection, melanoma checkpoint inhibitor therapy). This sensitivity of T-REX for extremely rare cells allowed the algorithm to reveal virus-specific CD4^+^ T cells without prior knowledge of their phenotype. T-REX should now be tested further to determine whether cells identified in SARS-CoV-2 also share a clonal origin. In addition, it is likely T-REX will be useful beyond immunology settings in paired comparisons of quantitative single cell data, such as discovery screening or paired analysis of tumor cells.

## Acknowledgements

We thank Monika Grabowska at Vanderbilt University for helpful work during a rotation on code that became part of the T-REX algorithm. Research was supported by the following funding resources: U01 AI125056 (S.M.B., A.G.A.P, L.M.M., R.B.T, J.A.W., and J.M.I.), R01 CA226833 (J.M.I., S.M.B.), U54 CA217450 (J.M.I.), T32 AI007496 (L.M.M.) and the Vanderbilt-Ingram Cancer Center (VICC, P30 CA68485).

## Author Contributions

S.M.B., A.G.A.P, L.M.M., J.A.W., and J.M.I. designed the data science study. A.G.A.P, L.M.M., J.A.L., W.W.K., R.B.T., and J.A.W. designed human rhinovirus challenge studies. A.G.A.P, L.M.M., and J.A.L. performed experimental work. S.M.B., A.G.A.P, L.M.M., J.A.L., J.A.W., and J.M.I. performed data analysis, developed figures, and wrote the manuscript. S.M.B., A.G.A.P, and L.M.M. compiled patient data. S.M.B and J.M.I. developed R scripts for data analysis and visualization. J.A.W. and J.M.I. provided financial support. All authors contributed in reviewing and editing the manuscript.

