## Supplemental Figures and Text for "Unsupervised machine learning reveals key immune cell subsets in COVID-19, rhinovirus infection, and cancer therapy"

### Supplemental figures and figure legends

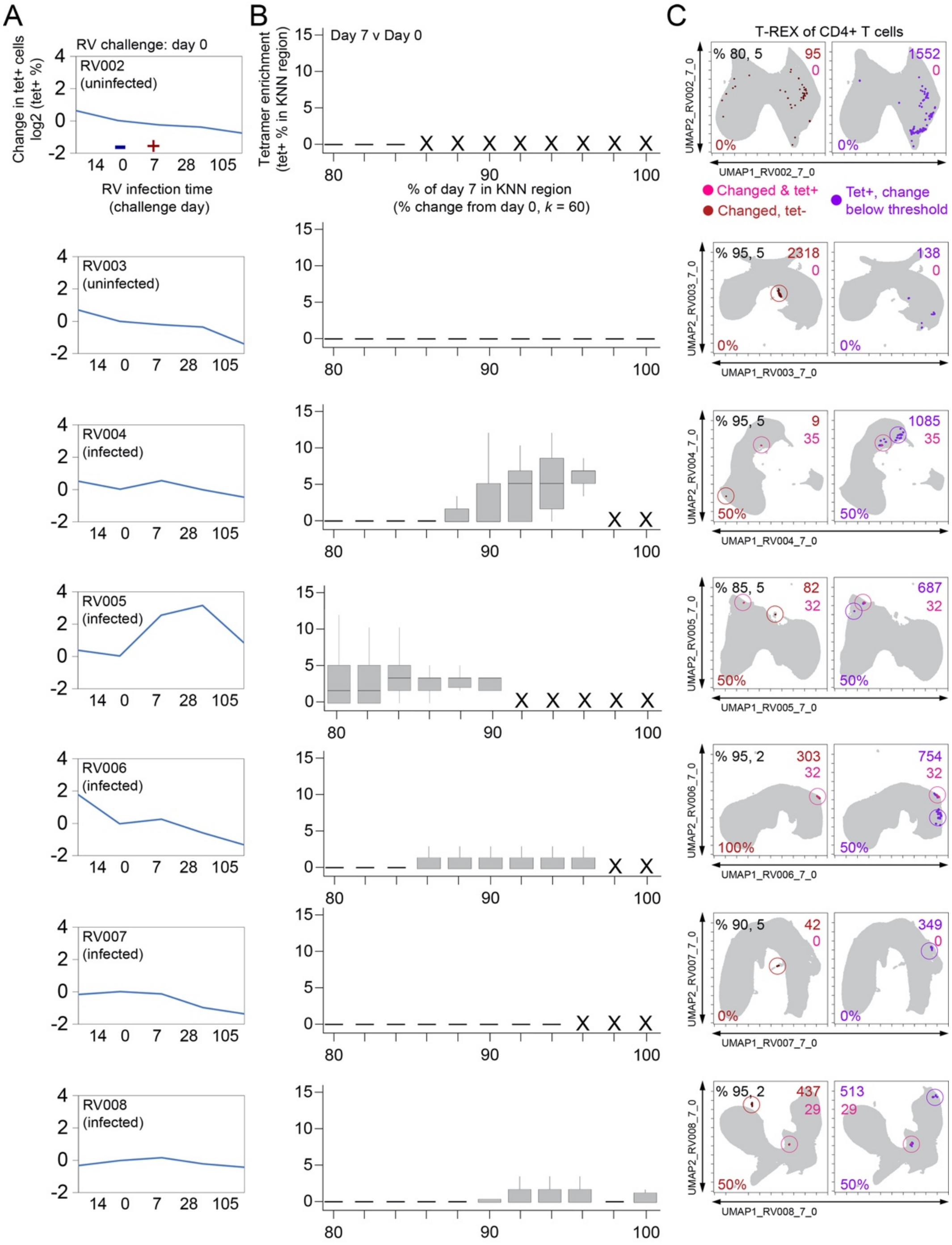


**Supplemental Figure 1 – T-REX identifies regions of great change enriched for tetramers in infected individuals.** Subjects RV002 through RV008 were experimentally infected with rhinovirus and CD4+ T cell signatures monitored by spectral flow cytometry in conjunction with tetramer staining during the course of infection. (A) Fold change in the number of tetramer-positive cells (log2) after rhinovirus challenge on day 0. (B) Box and whisker plots show KNN regions in terms of expansion during infection represented by percent change as well as percent of tetramer positive cells for day 0 and day. (C) UMAP plots for percent change and tetramer percent cutoff denoted in upper left corner in the left UMAP plot. Cell count in each region is in black as well as in the upper right of each UMAP plot for tetramer$^+$ regions changing (red), tetramer$^-$ regions changing (pink), and tetramer$^+$ regions with change below the expansion cutoff (purple). MEM labels are given for highly expanded and tetramer-enriched regions.

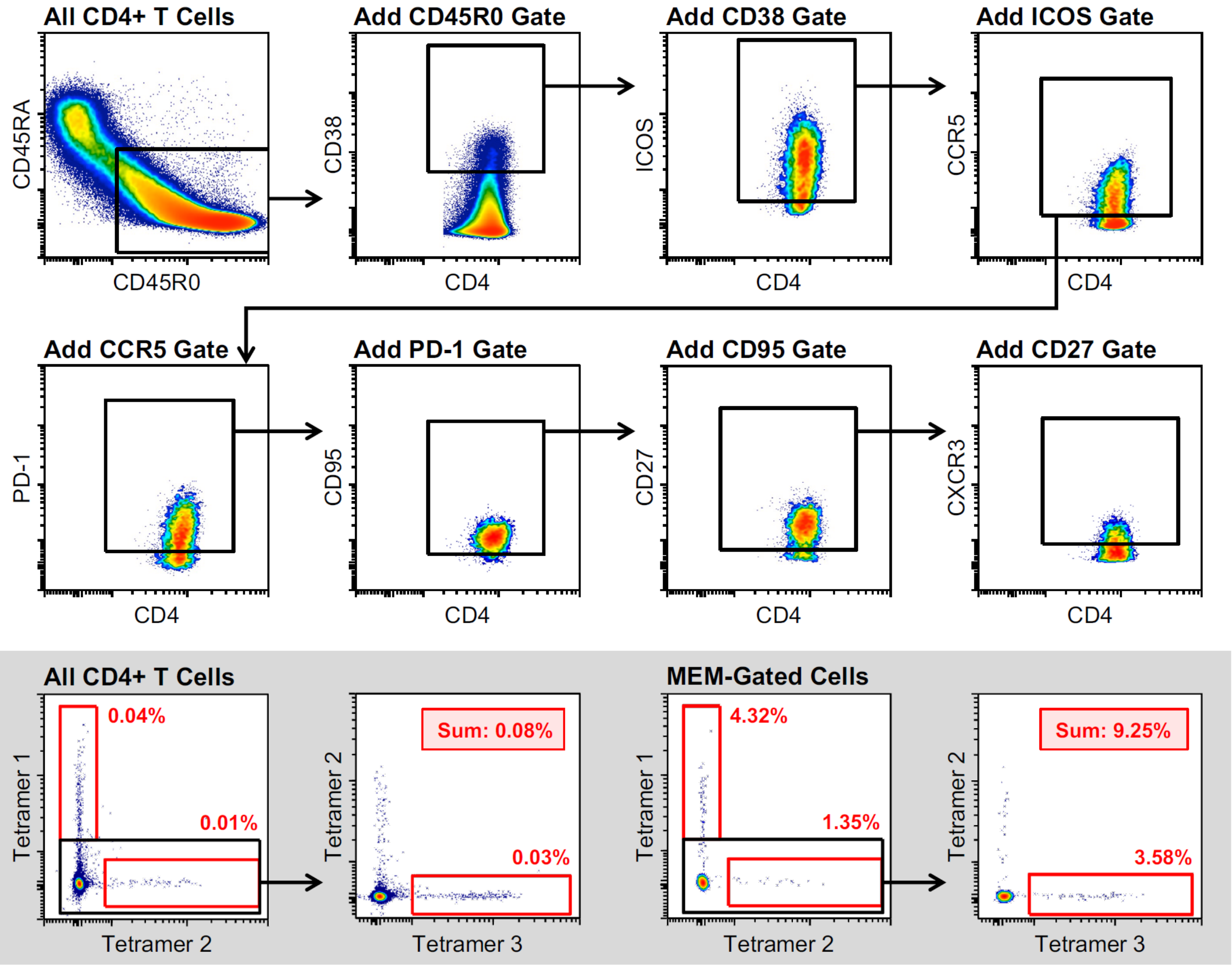


**Supplemental Figure 2 – MEM-derived gating strategy for the enrichment of rhinovirus-specific CD4+ T cells.** MEM-gated cells are derived from the combination of all depicted gates (CD45R0+ CD38+ ICOS+ CCR5+ PD-1+ CD95+ CD27+ CXCR3+. (Inset) Comparison of RV tetramer+ cell enrichment in ungated and MEM-gated cell populations.

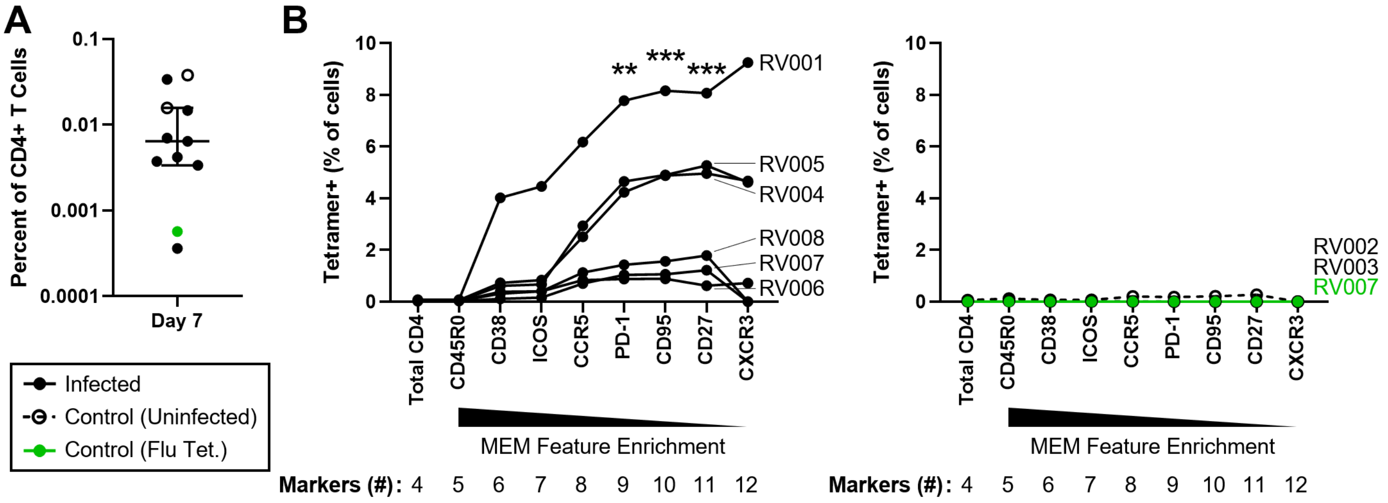


**Supplemental Figure 3 – T cell sorting strategy derived using T-REX effectively enriches for rhinovirus-specific cells in infected subjects**. (A) Precursor frequencies of total RV tetramer+ CD4+ T cells from all subjects on study day 7 (n=8 subjects). Median ± interquartile range. (B) Artificial sorting was performed using unenriched day 7 samples. Consensus MEM markers were individually added to the sorting strategy according to MEM feature enrichment, and corresponding total tetramer+ cell frequencies assessed. All cells were pre-gated for total CD4+ T cells (Live, Dump [CD14, CD19, CD8a]–, CD3+, CD4+). Two controls were utilized: rhinovirus tetramer staining was performed in subjects who remained uninfected following challenge as an infection control (n=2), and an unrelated influenza hemagglutinin tetramer was utilized in a rhinovirus-infected subject to confirm antigen specificity (n=1) (*right*). Tetramer enrichment with the addition of each marker was compared to the total CD4+ T cell population using Friedman's test with Dunn's multiple comparisons correction. **p ≤0.01; ***p ≤ 0.001.

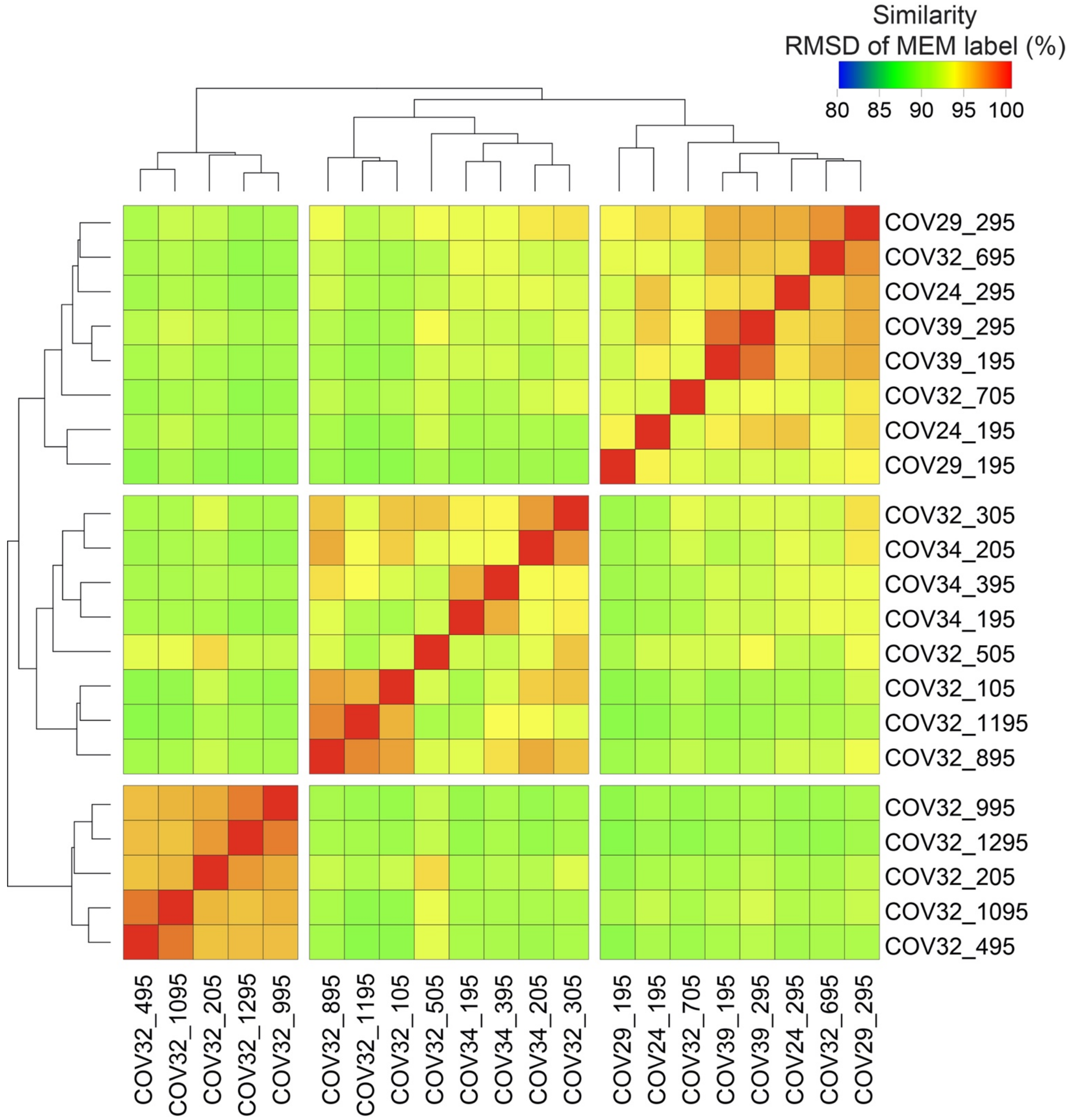


**Supplemental Figure 4 – RMSD on T-REX hotspot phenotypes from analysis of the COVID-19 CD4+ T cells identified three, distinct phenotypic groups** RMSD heatmap comparing MEM values for hotspots found by T-REX analysis on CD4+ T cells in COVID-19 samples.

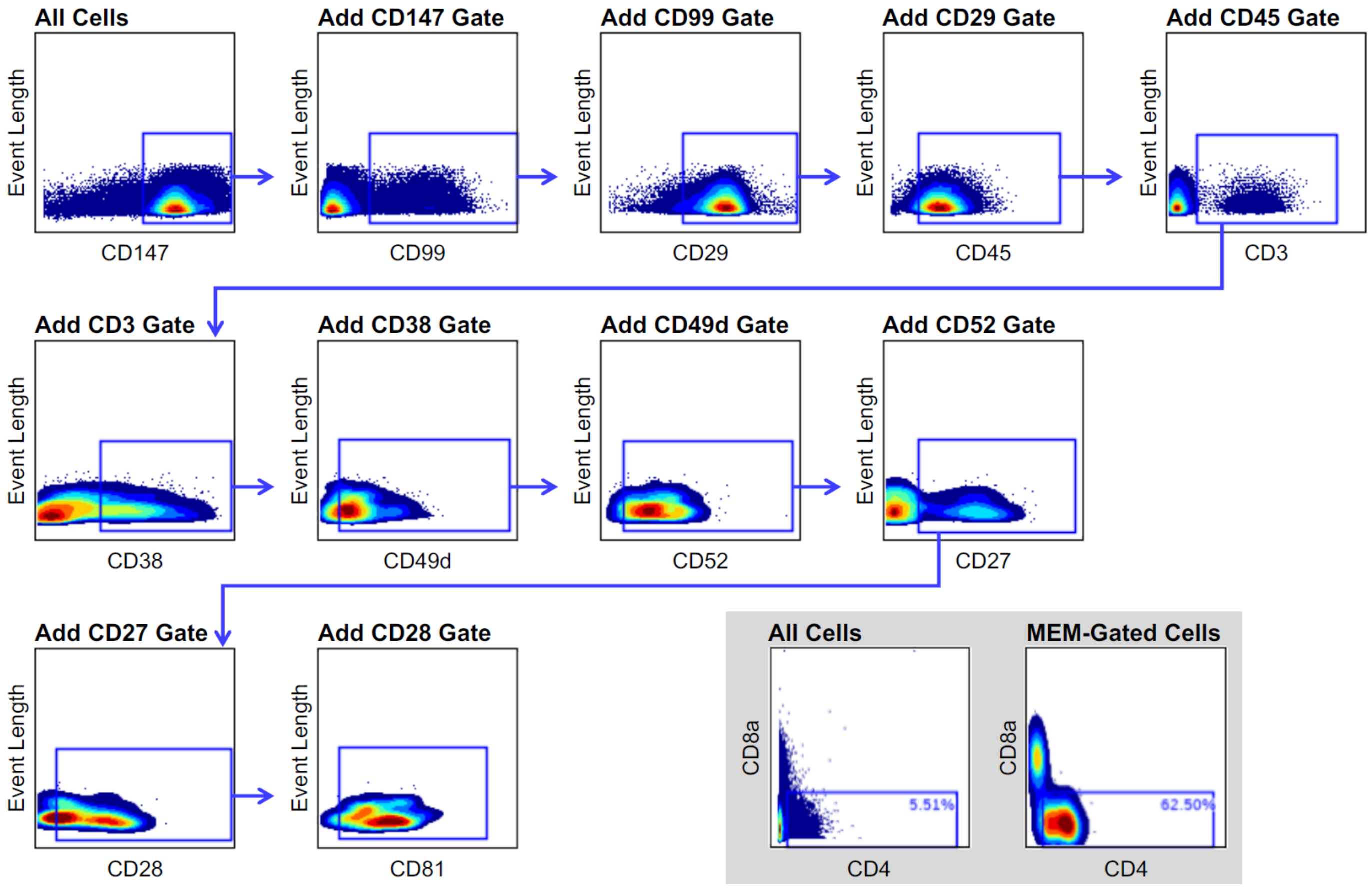


**Supplemental Figure 5 – MEM-derived gating strategy for the enrichment of CD4+ T cells in COVID-19 infected individuals.** MEM-gated cells are derived from the combination of all depicted gates (CD147+ CD99+ CD29+ CD45+ CD3+ CD38+ CD49d+ CD52+ CD27+ CD28+ CD81+). (Inset) Comparison of CD4+ T cell enrichment in ungated and MEM-gated cell populations.

### Supplemental Methods

#### Experimental Rhinovirus Infection Model

Healthy adult volunteers (ages 18-40) were enrolled in an experimental rhinovirus infection study. All subjects were judged to be seronegative to the challenge virus strain (RV-A16; serum neutralizing antibody titer ≤1:2). Subjects were inoculated with 100 $TCID_{50}$ of RV-A16 (FDA IND 15162) on study day 0, and were judged to be infected if they seroconverted to the challenge virus by study day 28 (≥4-fold increase in titer) and/or shed virus in nasal wash specimens during the first 5 days of infection, according to standard protocols [1]. Peripheral blood specimens were obtained sequentially over the course of infection to capture pre-infection immune fluctuations (days -14 and 0), the adaptive phase of infection (day 7), and convalescence (day 28). Peripheral blood mononuclear cells were isolated using density gradient centrifugation and viably cryopreserved for later analysis.

#### Rhinovirus Tetramer Staining and Flow Cytometry

Rhinovirus tetramer staining was performed as previously described [1-3]. Briefly, PBMCs were thawed, and all time points analyzed together in single experiments. Cells were stained with up to three unique rhinovirus MHCII tetramers, selected to match each individual's HLA type [3], and counterstained for viability and surface markers (Supplemental Table 3). An aliquot of tetramer-labeled cells was enriched using anti-PE magnetic beads for antigen-specific CD4+ T cell frequency calculation, as previously described [1-3]. Cells were fixed and permeabilized (True Nuclear Fixation and Permeabilization buffers; Biolegend, San Diego, CA, USA), and then stained for intracellular markers. Samples were analyzed using a 3-laser Aurora Northern Lights spectral flow cytometer (Cytek Biosciences, Fremont, CA, USA).
