## Supplemental Table 1 for "Unsupervised machine learning reveals key immune cell subsets in COVID-19, rhinovirus infection, and cancer therapy"

Supplemental Table 1 - Direction and degree of change for all samples as in Barone et al. Figure 6*

| Sample ID | Disease at day 0 | Sample source | Population analyzed | Total cells | Studied intervention | Outcome | Day compared | # cells, [0,5] | # cells, [5,15] | # cells, [15,85] | # cells, [85,95] | # cells, [95,100] | # cells, total | Change magnitude | Change direction |
|---|---|---|---|---|---|---|---|---|---|---|---|---|---|---|---|
| RV001 | Healthy | PBMC | CD4+ T cells | 1.37E+06 | Rhinovirus challenge | infected | 7 | 9 | 885 | 1363363 | 2080 | 1155 | 1367492 | 0.09% | 0.985 |
| RV002 | Healthy | PBMC | CD4+ T cells | 2.48E+06 | Rhinovirus challenge | uninfected | 7 | 9 | 329 | 2480954 | 0 | 0 | 2481292 | 0.00% | -1.000 |
| RV003 | Healthy | PBMC | CD4+ T cells | 2.62E+06 | Rhinovirus challenge | uninfected | 7 | 0 | 271 | 2613703 | 3615 | 1667 | 2619256 | 0.06% | 1.000 |
| RV004 | Healthy | PBMC | CD4+ T cells | 3.25E+06 | Rhinovirus challenge | infected | 7 | 291 | 4103 | 3243461 | 1316 | 11 | 3249182 | 0.01% | -0.927 |
| RV005 | Healthy | PBMC | CD4+ T cells | 2.30E+06 | Rhinovirus challenge | infected | 7 | 14 | 879 | 2303653 | 46 | 0 | 2304592 | 0.00% | -1.000 |
| RV006 | Healthy | PBMC | CD4+ T cells | 1.53E+06 | Rhinovirus challenge | infected | 7 | 0 | 280 | 1528931 | 1302 | 139 | 1530652 | 0.01% | 1.000 |
| RV007 | Healthy | PBMC | CD4+ T cells | 1.48E+06 | Rhinovirus challenge | infected | 7 | 6168 | 6858 | 1467017 | 95 | 0 | 1480138 | 0.42% | -1.000 |
| RV008 | Healthy | PBMC | CD4+ T cells | 1.16E+06 | Rhinovirus challenge | infected | 7 | 0 | 71 | 1154970 | 622 | 341 | 1156004 | 0.03% | 1.000 |
| COV24 | COVID-19 | PBMC | All cells | 6.34E+04 | N/A | ICU | 3 | 3 | 1105 | 58091 | 3580 | 647 | 63426 | 1.025% | 0.991 |
| COV26 | COVID-19 | PBMC | All cells | 1.19E+05 | N/A | ICU | 6 | 4 | 338 | 111308 | 5389 | 2409 | 119448 | 2.020% | 0.997 |
| COV27 | COVID-19 | PBMC | All cells | 4.90E+04 | N/A | ICU | 1 | 12 | 1859 | 45592 | 1532 | 11 | 49006 | 0.047% | -0.043 |
| COV29 | COVID-19 | PBMC | All cells | 1.64E+05 | N/A | ICU | 4 | 12208 | 17659 | 113306 | 17287 | 3920 | 164380 | 9.811% | -0.514 |
| COV32 | COVID-19 | PBMC | All cells | 3.21E+05 | N/A | non ICU | 6 | 119305 | 18635 | 51651 | 31496 | 99643 | 320730 | 68.266% | -0.090 |
| COV34 | COVID-19 | PBMC | All cells | 4.97E+05 | N/A | ICU | 7 | 97380 | 70135 | 164237 | 99849 | 64921 | 496522 | 32.688% | -0.200 |
| COV35 | COVID-19 | PBMC | All cells | 8.14E+04 | N/A | ICU | 4 | 141 | 3044 | 74262 | 3341 | 610 | 81398 | 0.923% | 0.625 |
| COV36 | COVID-19 | PBMC | All cells | 2.96E+05 | N/A | non ICU | 5 | 5494 | 5497 | 271632 | 7703 | 6112 | 296438 | 3.915% | 0.053 |
| COV37 | COVID-19 | PBMC | All cells | 5.81E+03 | N/A | ICU | 5 | 1839 | 523 | 977 | 538 | 1937 | 5814 | 64.947% | 0.026 |
| COV39 | COVID-19 | PBMC | All cells | 1.94E+05 | N/A | ICU | 4 | 45209 | 12696 | 88567 | 30561 | 16717 | 193750 | 31.962% | -0.460 |
| COV40 | COVID-19 | PBMC | All cells | 4.84E+04 | N/A | ICU | 6 | 347 | 1477 | 42432 | 3151 | 959 | 48366 | 2.700% | 0.469 |
| COV41 | COVID-19 | PBMC | All cells | 9.82E+04 | N/A | ICU | 4 | 15488 | 9468 | 62216 | 8458 | 2598 | 98228 | 18.412% | -0.713 |
| MB004 | Melanoma | PBMC | All cells | 1.69E+05 | α-PD immunotherapy | complete | 3 | 4975 | 20025 | 115912 | 22236 | 5672 | 168820 | 6.307% | 0.065 |
| MB005 | Melanoma | PBMC | All cells | 1.54E+05 | α-PD immunotherapy | complete | 3 | 1117 | 1636 | 149069 | 2042 | 60 | 153924 | 0.765% | -0.898 |
| MB006 | Melanoma | PBMC | All cells | 1.92E+05 | α-PD immunotherapy | progressive | 5 | 11418 | 4735 | 172333 | 3476 | 276 | 192238 | 6.083% | -0.953 |
| MB007 | Melanoma | PBMC | All cells | 1.72E+05 | α-PD immunotherapy | progressive | 3 | 536 | 2916 | 168874 | 154 | 0 | 172480 | 0.311% | -1.000 |
| MB008 | Melanoma | PBMC | All cells | 2.33E+05 | α-PD immunotherapy | partial | 3 | 5991 | 7137 | 211481 | 8094 | 491 | 233194 | 2.780% | -0.849 |
| MB009 | Melanoma | PBMC | All cells | 4.65E+05 | α-PD immunotherapy | stable | 3 | 1838 | 6903 | 455082 | 1401 | 0 | 465224 | 0.395% | -1.000 |
| MB010 | Melanoma | PBMC | All cells | 3.94E+05 | α-PD immunotherapy | partial | 3 | 0 | 478 | 392391 | 851 | 20 | 393740 | 0.005% | 1.000 |
| MB012 | Melanoma | PBMC | All cells | 1.65E+05 | α-PD immunotherapy | partial | 3 | 459 | 6870 | 147120 | 7973 | 2306 | 164728 | 1.679% | 0.668 |
| MB013 | Melanoma | PBMC | All cells | 4.29E+05 | α-PD immunotherapy | partial | 6 | 56 | 579 | 424404 | 3654 | 449 | 429142 | 0.118% | 0.778 |
| MB041 | Melanoma | PBMC | All cells | 3.45E+05 | α-PD immunotherapy | progressive | 3 | 0 | 0 | 344611 | 13 | 0 | 344624 | 0.000% | 0.000 |
| AML001 | AML | PBMC | All cells | 6.31E+04 | Chemotherapy | CR | 5 | 31498 | 7 | 131 | 161 | 31341 | 63138 | 99.526% | -0.002 |
| AML002 | AML | PBMC | All cells | 7.40E+04 | Chemotherapy | CR | 8 | 29663 | 5074 | 5134 | 3878 | 30209 | 73958 | 80.954% | 0.009 |
| AML003 | AML | PBMC | All cells | 6.83E+05 | Chemotherapy | PD | 5 | 1667 | 48603 | 556652 | 55640 | 20082 | 682644 | 3.186% | 0.847 |
| AML004 | AML | PBMC | All cells | 3.41E+05 | Chemotherapy | CR | 8 | 139293 | 21982 | 25628 | 16694 | 137025 | 340622 | 81.122% | -0.008 |
| AML007 | AML | PBMC | All cells | 1.94E+05 | Chemotherapy | CR | 8 | 6203 | 14309 | 144865 | 22967 | 5370 | 193714 | 5.974% | -0.072 |

*For studied interventions and outcomes, see original manuscripts for detailed information. Day compared indicates the actual day a sample was taken relative to a day 0 comparison in all cases. #cells indicates the number of cells in each percentile grouping (e.g. [0,5] indicates the number of cells whose KNN area had less than or equal to 5% change in T-REX analysis). Change magnitude and direction are as in Figure 6A.
