## Supplemental Table 2a for "Unsupervised machine learning reveals key immune cell subsets in COVID-19, rhinovirus infection, and cancer therapy"

**Supplemental Table 2a - MEM Labels for Enriched Features in All Cell Hotspots from COVID-19 Patients in Dataset 2**

| patient_id | degree_change | direction_change | DBSCAN_cluster | total_counts | Marker Enrichment Modeling (MEM) Label |
|---|---|---|---|---|---|
| COV24 | 1.025 | 0.991 | 1 | 85 | 2401095 : ▲ CD147+6 CD99+6 CD29+5 CD38+4 CD3+3 CD49d+2 CD81+2 CD52+2 CD27+2 CD28+2 CD55+1 CD62L+1 CD56+1 CD5+1 CD4+1 ▼ None |
| | | | 2 | 174 | 2402095 : ▲ CD9+5 CD29+5 CD147+4 CD99+4 IgD+1 CD55+1 CX3CR1+1 CD34+1 ▼ None |
| | | | 3 | 157 | 2403095 : ▲ CD147+5 CD9+4 CD29+3 CD81+2 Siglec-8+2 CD39+2 CD55+1 CD123+1 CD24+1 CD99+1 CD8a+1 ▼ None |
| | | | 4 | 54 | 2404095 : ▲ CD147+6 CD99+6 CD29+4 CD38+4 CD8a+4 CD49d+2 CD3+2 CD27+2 CD39+2 CD81+1 CD52+1 CD55+1 CD56+1 ▼ None |
| | | | 5 | 135 | 2405095 : ▲ CD147+6 CD29+5 CD56+5 CD99+5 CD38+4 CD49d+2 CD81+2 CD7+2 CD52+1 CD55+1 CD8a+1 ▼ None |
| COV26 | 2.020 | 0.997 | 1 | 1062 | 2601095 : ▲ CD147+6 CD99+5 CD29+4 CD38+3 CD55+2 CD14+2 CD64+1 CD56+1 CD39+1 ▼ None |
| | | | 2 | 974 | 2602095 : ▲ CD9+4 CD147+4 CD81+1 CD55+1 Siglec-8+1 CD29+1 CD8a+1 ▼ None |
| | | | 3 | 163 | 2603095 : ▲ CD9+7 CD29+6 CD147+5 CD99+5 CX3CR1+4 CD34+2 IgD+1 CD43+1 CD45RB+1 CD55+1 CD20+1 CD24+1 CD56+1 ▼ None |
| | | | 4 | 58 | 2604095 : ▲ CD147+6 CD29+3 CD55+2 CD99+2 CD49d+1 CD52+1 CD56+1 ▼ None |
| | | | 5 | 97 | 2605095 : ▲ CD38+5 CD147+5 CD123+4 CD55+2 CD99+2 CD9+1 CD62L+1 CD8a+1 ▼ None |
| COV27 | 0.047 | -0.043 | 1 | 8 | 270105 : ▲ CD147+7 CD99+6 CD29+4 CD64+3 CD38+3 CD55+2 CD14+2 CD56+2 CD62L+1 CD39+1 CD8a+1 ▼ None |
| | | | 2 | 4 | 270205 : ▲ CD99+6 CD57+4 CD147+4 CD45RA+3 CD56+3 CD29+2 CD38+2 CD43+1 CD81+1 CD8a+1 ▼ None |
| | | | 3 | 8 | 2703095 : ▲ CD147+4 CD16+3 CD24+2 CD55+1 CD29+1 CD38+1 CD39+1 ▼ None |
| | | | 4 | 2 | 2704095 : ▲ CD147+4 CD16+2 CD55+1 CD24+1 CD56+1 CD99+1 CD39+1 CD8a+1 ▼ None |
| COV29 | 9.811 | -0.514 | 1 | 11879 | 290105 : ▲ CD16+5 CD147+4 CD55+3 CD24+3 CD29+2 CD127+2 CD39+2 CD45+1 CD38+1 CD8a+1 ▼ None |
| | | | 2 | 284 | 290205 : ▲ CD147+6 CD16+6 CD55+4 CD24+4 CD29+3 CD127+3 CD39+3 CD38+2 CD45+1 CD33+1 CD11c+1 CD43+1 CD64+1 CD62L+1 CD56+1 CD99+1 CD8a+1 ▼ None |
| | | | 3 | 2495 | 2903095 : ▲ CD147+4 CD55+2 CD29+2 CD24+2 CD39+2 CD16+2 CD38+1 ▼ None |
| | | | 4 | 192 | 2904095 : ▲ CD147+4 CD29+2 CD64+1 CD24+1 CD39+1 CD8a+1 ▼ None |
| | | | 5 | 759 | 2905095 : ▲ CD9+4 CD147+4 Siglec-8+3 CD81+2 CD29+2 CD39+2 CD45+1 CD55+1 CD34+1 CD127+1 CD24+1 CD38+1 CD99+1 CD8a+1 ▼ None |
| | | | 6 | 246 | 2906095 : ▲ CD147+6 CD9+4 CD55+3 Siglec-8+3 CD29+3 CD24+3 CD39+3 CD16+3 CD81+2 CD38+2 CD45+1 CD11c+1 CD43+1 CD34+1 CD62L+1 CD127+1 CD56+1 CD99+1 CD8a+1 ▼ None |
| COV32 | 68.266 | -0.090 | 1 | 99147 | 320105 : ▲ CD147+5 CD55+3 CD16+3 CD29+2 CD24+2 CD39+2 CD62L+1 CD38+1 CD56+1 ▼ None |
| | | | 2 | 5835 | 320205 : ▲ CD147+6 CD55+4 CD24+4 CD16+4 CD29+3 CD39+3 CD62L+2 CD45+1 CD11c+1 CD43+1 CD64+1 CD38+1 CD56+1 CD99+1 CD8a+1 ▼ None |
| | | | 3 | 165 | 320305 : ▲ CD147+5 CD55+3 CD29+3 CD16+3 CD9+2 CD24+2 CD39+2 CD62L+1 CD38+1 CD56+1 CD99+1 CD8a+1 ▼ None |
| | | | 4 | 10719 | 320405 : ▲ CD147+6 CD29+5 CD55+4 CD38+4 CD99+4 CD64+3 CD14+3 CD45+2 CD11c+2 CD62L+2 CD56+2 CD39+2 CD33+1 HLA-DR+1 CD49d+1 CD137+1 CD141+1 CD85j+1 ▼ None |
| | | | 5 | 765 | 320505 : ▲ CD123+8 Siglec-8+8 CD127+8 CD28+8 CD43+5 CD147+5 CD55+3 CD39+3 CD16+3 CD20+2 CD29+2 CD24+2 CD62L+1 CD38+1 CD56+1 ▼ None |
| | | | 6 | 937 | 320605 : ▲ CD147+7 CD55+5 CD29+5 CD38+5 CD99+4 CD16+4 CD64+3 CD14+3 CD24+3 CD39+3 CD45+2 CD11c+2 CD62L+2 CD56+2 CD33+1 HLA-DR+1 CD49d+1 CD43+1 CD52+1 CD137+1 CD141+1 CD85j+1 CD8a+1 ▼ None |
| | | | 7 | 1014 | 320705 : ▲ CD147+5 CD9+4 Siglec-8+3 CD81+2 CD29+2 CD24+2 CD39+2 CD45+1 CD49d+1 CD55+1 CD34+1 CD62L+1 CD127+1 CD38+1 CD56+1 CD99+1 CD8a+1 ▼ None |
| | | | 8 | 435 | 320805 : ▲ CD57+4 CD3+4 CD45RA+4 CD29+3 CD38+3 CD147+3 CD99+3 CD45+2 CD43+2 CD81+2 CD56+2 CD8a+2 CD11c+1 CD49d+1 CD52+1 CD161+1 ▼ None |
| | | | 9 | 116 | 320905 : ▲ CD57+5 CD29+4 CD3+3 CD147+3 CD99+3 CD45+2 CD43+2 CD81+2 CD52+2 CD49d+1 CD38+1 CD8a+1 CD5+1 ▼ None |
| | | | 2 | 12 | 3202095 : ▲ CD147+7 CD24+5 CD16+5 CD55+4 CD29+4 CD39+4 CD45+2 CD11c+2 CD38+2 CD56+2 CD43+1 CD64+1 CD9+1 CD62L+1 CD127+1 CD99+1 CD8a+1 ▼ None |
| | | | 3 | 76142 | 3203095 : ▲ CD147+4 CD24+3 CD16+3 CD55+2 CD29+2 CD39+2 CD45+1 CD11c+1 CD38+1 ▼ None |
| | | | 5 | 493 | 3205095 : ▲ CD123+8 Siglec-8+8 CD127+8 CD28+8 CD43+5 CD147+4 CD20+3 CD24+3 CD39+3 CD16+3 CD55+2 CD29+2 CD45+1 CD11c+1 CD9+1 ▼ None |
| | | | 6 | 1802 | 3206095 : ▲ CD147+7 CD29+5 CD99+5 CD55+4 CD38+4 CD16+4 CD11c+3 CD9+3 CD14+3 CD24+3 CD39+3 CD45+2 CD64+2 CD56+2 CD33+1 HLA-DR+1 CD49d+1 CD52+1 CD137+1 CD62L+1 CD141+1 CD85j+1 CD8a+1 ▼ None |
| | | | 7 | 1307 | 3207095 : ▲ CD9+5 CD147+5 CD81+3 Siglec-8+3 CD29+3 CD24+3 CD45+2 CD55+2 CD99+2 CD39+2 CD11c+1 CD52+1 CD34+1 CD38+1 CD56+1 CD8a+1 ▼ None |
| | | | 8 | 572 | 3208095 : ▲ CD57+5 CD45RA+5 CD99+5 CD3+4 CD147+4 CD29+3 CD56+3 CD8a+3 CD45+2 CD43+2 CD81+2 CD38+2 CD11c+1 CD49d+1 CD52+1 CD161+1 CD7+1 ▼ None |
| | | | 9 | 1029 | 3209095 : ▲ CD57+5 CD99+5 CD29+4 CD43+3 CD3+3 CD147+3 CD45+2 CD49d+2 CD81+2 CD52+2 CD45RB+1 CD45RA+1 CD56+1 CD8a+1 CD5+1 CD4+1 ▼ None |
| | | | 10 | 294 | 32010095 : ▲ CD29+5 CD147+5 CD38+4 CD56+4 CD99+4 CD45+2 CD49d+2 CD81+2 CD52+2 CD43+1 CD55+1 CD161+1 CD137+1 CD62L+1 CD45RA+1 CD8a+1 CD7+1 ▼ None |
| | | | 11 | 12291 | 32011095 : ▲ CD147+6 CD29+5 CD99+4 CD11c+3 CD55+3 CD9+3 CD14+3 CD38+3 CD45+2 CD64+2 CD56+2 CD39+2 CD33+1 CD137+1 CD62L+1 CD85j+1 ▼ None |
| | | | 12 | 2093 | 32012095 : ▲ CD147+6 CD24+4 CD16+4 CD55+3 CD29+3 CD39+3 CD45+2 CD11c+2 CD43+1 CD9+1 CD38+1 CD56+1 CD99+1 CD8a+1 ▼ None |
| | | | 13 | 471 | 32013095 : ▲ CD29+5 CD147+5 CD99+5 CD38+4 CD8a+4 CD45+3 CD49d+3 CD3+3 CD52+3 CD81+2 CD27+2 HLA-DR+1 CD43+1 CD137+1 CD56+1 CD39+1 ▼ None |
| | | | 14 | 2290 | 32014095 : ▲ CD9+10 CD29+8 CD147+7 CD99+6 CD34+5 CD55+4 CD20+4 CD137+3 CD56+3 CD81+1 CX3CR1+1 CD14+1 CD161+1 CD62L+1 CD38+1 ▼ None |
| | | | 15 | 273 | 32015095 : ▲ CD147+6 CD99+6 CD29+5 CD38+4 CD45+3 CD3+3 CD52+3 CD27+3 CD49d+2 CD28+2 CD43+1 CD81+1 CD55+1 CD137+1 CD62L+1 CD56+1 CD5+1 CD4+1 ▼ None |
| COV34 | 32.688 | -0.200 | 1 | 92331 | 340105 : ▲ CD147+4 CD55+3 CD24+3 CD16+3 CD29+2 CD62L+1 CD127+1 CD56+1 CD39+1 CD8a+1 ▼ None |
| | | | 2 | 370 | 340205 : ▲ CD147+5 CD16+4 CD55+3 CD29+3 CD99+2 CD45+1 CD11c+1 CD49d+1 CD43+1 CD123+1 CD85j+1 CD56+1 CD8a+1 ▼ None |
| | | | 3 | 935 | 340305 : ▲ CD24+5 CD147+5 CD55+4 CD16+4 CD29+2 CD45+1 CD43+1 CD62L+1 CD38+1 CD56+1 CD39+1 CD8a+1 ▼ None |
| | | | 4 | 1762 | 340405 : ▲ CD147+6 CD29+5 CD55+4 CD38+4 CD99+4 CD64+3 CD14+3 CD62L+3 CD56+2 CD39+2 CD45+1 CD11c+1 CD137+1 CD141+1 CD85j+1 CD8a+1 ▼ None |
| | | | 5 | 340 | 340505 : ▲ CD147+5 CD55+4 CD24+4 CD16+4 CD29+3 CD127+2 CD39+2 CD8a+2 CD45+1 CD43+1 CD64+1 CD62L+1 CD38+1 CD56+1 CD99+1 ▼ None |
| | | | 7 | 704 | 340705 : ▲ CD147+4 CD55+2 CD29+2 CD24+1 CD56+1 CD99+1 CD39+1 CD8a+1 CD16+1 ▼ None |
| | | | 8 | 907 | 340805 : ▲ CD38+4 CD99+4 CD29+3 CD45RA+3 CD147+3 CD57+2 CD56+2 CD45+1 CD43+1 CD81+1 CD8a+1 ▼ None |
| | | | 9 | 11 | 340905 : ▲ CD57+4 CD99+4 CD29+3 CD147+3 CD8a+3 CD38+2 CD45+1 CD49d+1 CD43+1 CD3+1 CD81+1 CD52+1 CD45RA+1 CD5+1 ▼ None |
| | | | 3 | 177 | 3403095 : ▲ CD147+5 CD24+4 CD16+4 CD55+3 CD29+3 CD39+2 CD45+1 CD43+1 CD64+1 CD62L+1 CD38+1 CD56+1 CD8a+1 ▼ None |
| | | | 6 | 59047 | 3406095 : ▲ CD147+4 CD16+4 CD55+3 CD29+2 CD24+2 CD62L+1 CD38+1 CD56+1 CD39+1 CD8a+1 ▼ None |
| | | | 8 | 299 | 3408095 : ▲ CD45RA+5 CD99+5 CD57+4 CD29+4 CD38+4 CD147+4 CD56+4 CD43+2 CD8a+2 CD45+1 CD49d+1 CD81+1 CD161+1 CD7+1 ▼ None |
| | | | 9 | 786 | 3409095 : ▲ CD57+5 CD99+5 CD29+4 CD8a+4 CD147+3 CD45+2 CD43+2 CD3+2 CD81+2 CD45RA+2 CD49d+1 CD52+1 CD5+1 ▼ None |
| | | | 10 | 893 | 34010095 : ▲ CD9+4 CD29+4 CD55+3 CD147+3 CD99+3 CD43+1 CD64+1 CX3CR1+1 CD34+1 CD24+1 CD56+1 ▼ None |
| | | | 11 | 941 | 34011095 : ▲ CD9+4 CD147+4 CD29+3 CD81+2 CD55+2 Siglec-8+2 CD8a+2 CD45+1 CD34+1 CD127+1 CD24+1 CD38+1 CD56+1 CD99+1 CD39+1 ▼ None |
| | | | 12 | 1338 | 34012095 : ▲ CD147+6 CD29+5 CD99+5 CD55+3 CD38+3 CD62L+2 CD56+2 CD45+1 CD64+1 CD14+1 CD137+1 CD141+1 CD39+1 CD8a+1 ▼ None |
| | | | 13 | 312 | 34013095 : ▲ CD147+5 CD55+3 CD29+3 CD62L+2 CD24+2 CD16+2 CD38+1 CD56+1 CD99+1 CD39+1 CD8a+1 ▼ None |
| | | | 14 | 207 | 34014095 : ▲ CD147+7 CD29+6 CD38+6 CD45RA+5 CD99+4 CD39+4 CD49d+3 CD43+2 CD81+2 CD62L+2 CD27+2 CD56+2 CD45+1 HLA-DR+1 CD55+1 CD137+1 CD85j+1 CD8a+1 ▼ None |
| | | | 15 | 247 | 34015095 : ▲ CD147+5 CD29+3 CD24+2 CD8a+2 CD43+1 CD55+1 CD64+1 CD38+1 CD56+1 CD99+1 CD39+1 CD16+1 ▼ None |
| | | | 16 | 342 | 34016095 : ▲ CD147+5 CD16+5 CD55+4 CD29+4 CD62L+3 CD24+3 CD39+2 CD8a+2 CD45+1 CD43+1 CD64+1 CD38+1 CD56+1 CD99+1 ▼ None |
| COV35 | 0.923 | 0.625 | 1 | 36 | 350105 : ▲ CD147+5 CD55+3 CD62L+1 CD24+1 CD16+1 ▼ None |
| | | | 2 | 92 | 350205 : ▲ CD99+5 CD45RA+4 CD57+3 CD147+2 CD56+2 CD43+1 CD29+1 CD38+1 CD8a+1 ▼ None |
| | | | 3 | 204 | 3503095 : ▲ CD9+5 CD147+5 CD81+2 CD55+1 Siglec-8+1 CD29+1 CD127+1 CD24+1 CD39+1 CD8a+1 ▼ None |
| | | | 4 | 202 | 3504095 : ▲ CD9+5 CD29+5 CD147+3 CD99+2 CX3CR1+1 ▼ None |
| | | | 5 | 194 | 3505095 : ▲ CD9+7 CD29+6 CD147+5 CD99+4 CD55+2 CD34+2 CX3CR1+1 CD38+1 CD56+1 ▼ None |
| COV36 | 3.915 | 0.053 | 1 | 2378 | 360105 : ▲ CD147+6 CD29+5 CD55+4 CD38+4 CD14+3 CD99+3 CD45+2 CD11c+2 CD64+2 CD33+1 HLA-DR+1 CD62L+1 CD56+1 CD39+1 ▼ None |
| | | | 2 | 2310 | 360205 : ▲ CD9+3 CD29+3 IgD+1 CX3CR1+1 CD24+1 CD147+1 CD99+1 ▼ None |
| | | | 3 | 352 | 360305 : ▲ CD55+4 CD147+4 CD24+2 CD45+1 CD64+1 CD29+1 CD62L+1 CD39+1 CD16+1 ▼ None |
| | | | 4 | 421 | 360405 : ▲ CD147+4 CD55+3 CD24+3 CD16+3 CD127+2 CD45+1 CD29+1 CD39+1 ▼ None |
| | | | 1 | 3816 | 3601095 : ▲ CD147+6 CD29+5 CD99+5 CD38+3 CD45+2 CD11c+2 CD55+2 CD14+2 CD39+2 CD33+1 HLA-DR+1 CD64+1 CD9+1 CD56+1 ▼ None |
| | | | 5 | 2152 | 3605095 : ▲ CD9+9 CD29+8 CD147+7 CD99+6 CD34+5 CD55+3 CD20+3 CD137+2 CD56+2 CD161+1 CD38+1 ▼ None |
| COV37 | 64.947 | 0.026 | 1 | 1410 | 370105 : ▲ CD147+4 CD55+2 CD24+1 CD16+1 ▼ None |
| | | | 2 | 229 | 370205 : ▲ CD99+4 CD3+2 CD52+1 CD29+1 CD27+1 CD147+1 ▼ None |
| | | | 3 | 113 | 370305 : ▲ CD38+5 CD147+3 CD99+3 CD29+1 CD45RA+1 CD56+1 ▼ None |
| | | | 4 | 77 | 370405 : ▲ CD147+6 CD55+4 CD29+3 CD38+3 CD64+2 CD99+2 CD14+1 CD56+1 CD39+1 ▼ None |
| | | | 5 | 1639 | 3705095 : ▲ CD147+4 CD9+3 CD81+1 Siglec-8+1 CD29+1 CD127+1 CD39+1 CD8a+1 ▼ None |
| | | | 6 | 254 | 3706095 : ▲ CD9+10 CD29+9 CD147+8 CD99+7 CD34+6 CD55+5 CD20+4 CD56+4 CD137+3 CD38+2 CX3CR1+1 ▼ None |
| COV39 | 31.962 | -0.460 | 1 | 24346 | 390105 : ▲ CD147+5 CD16+5 CD55+4 CD45+1 CD64+1 CD29+1 CD62L+1 CD24+1 CD39+1 ▼ None |
| | | | 2 | 17959 | 390205 : ▲ CD9+4 CD29+4 CD147+2 IgD+1 CD99+1 ▼ None |
| | | | 4 | 1198 | 390405 : ▲ CD29+5 CD147+5 CD55+4 CD9+4 CD16+4 CD24+2 CD45+1 CD11c+1 IgD+1 CD64+1 CX3CR1+1 CD62L+1 CD38+1 CD99+1 CD39+1 ▼ None |
| | | | 5 | 1173 | 390505 : ▲ CD147+7 CD99+5 CD29+4 CD38+4 CD55+3 CD64+3 CD45+1 CD11c+1 CD14+1 CD62L+1 CD56+1 CD39+1 ▼ None |
| | | | 10 | 194 | 39010095 : ▲ CD99+6 CD29+5 CD147+5 CD45+3 CD27+3 CD3+2 CD52+2 CD38+2 CD28+2 CD49d+1 CD5+1 ▼ None |
| | | | 11 | 222 | 39011095 : ▲ CD9+5 CD147+3 CD29+2 CD45+1 CD81+1 Siglec-8+1 CD39+1 ▼ None |
| | | | 11 | 7 | 3901105 : ▲ CD147+4 CD9+3 CD45+1 CD81+1 CD55+1 Siglec-8+1 CD29+1 CD24+1 CD39+1 ▼ None |
| | | | 3 | 9986 | 3903095 : ▲ CD147+4 CD55+1 CD29+1 CD24+1 CD16+1 ▼ None |
| | | | 6 | 1930 | 3906095 : ▲ CD147+4 CD29+1 ▼ None |
| | | | 7 | 2165 | 3907095 : ▲ CD147+6 CD29+4 CD11c+3 CD45+2 CD55+2 CD99+2 HLA-DR+1 CD49d+1 CD64+1 CD14+1 CD38+1 CD56+1 CD39+1 ▼ None |
| | | | 8 | 993 | 3908095 : ▲ CD147+5 CD24+3 CD16+3 CD55+2 CD29+2 CD45+1 CD38+1 CD39+1 ▼ None |
| | | | 9 | 722 | 3909095 : ▲ CD29+5 CD99+5 CD147+4 CD38+3 CD45+2 CD8a+2 CD49d+1 CD3+1 CD52+1 CD27+1 ▼ None |
| COV40 | 2.700 | 0.469 | 1 | 228 | 400105 : ▲ CD147+6 CD55+4 CD29+4 CD38+4 CD99+3 CD64+2 CD62L+2 CD45+1 ▼ None |
| | | | 2 | 52 | 400205 : ▲ CD147+8 CD38+7 CD29+6 CD39+6 CD45RA+5 CD99+5 CD27+4 CD45+2 HLA-DR+1 CD49d+1 CD81+1 CD62L+1 ▼ None |
| | | | 3 | 748 | 4003095 : ▲ CX3CR1+6 CD9+6 CD29+6 CD147+3 IgD+2 CD99+2 CD33+1 CD11c+1 HLA-DR+1 CD24+1 ▼ None |
| | | | 4 | 57 | 4004095 : ▲ CX3CR1+5 CD9+4 CD29+3 IgD+2 CD11c+1 CD147+1 ▼ None |
| | | | 5 | 52 | 4005095 : ▲ CX3CR1+4 IgD+2 CD9+2 CD11c+1 CD29+1 ▼ None |
| COV41 | 18.412 | -0.713 | 1 | 10355 | 410105 : ▲ CD147+5 CD16+5 CD55+3 CD24+2 CD45+1 CD64+1 CD29+1 CD62L+1 CD39+1 ▼ None |
| | | | 2 | 3930 | 410205 : ▲ CD9+2 CD29+2 IgD+1 CX3CR1+1 CD147+1 ▼ None |
| | | | 3 | 493 | 410305 : ▲ CD24+3 CD147+3 CD55+2 CD29+1 CD62L+1 CD39+1 CD16+1 ▼ None |
| | | | 4 | 404 | 410405 : ▲ CD147+5 CD55+3 CD24+2 CD16+2 CD64+1 CD29+1 CD39+1 ▼ None |
| | | | 5 | 211 | 410505 : ▲ CD147+6 CD29+5 CD99+5 CD55+3 CD64+3 CD38+3 CD45+1 CD14+1 CD56+1 CD39+1 ▼ None |
| | | | 6 | 70 | 410605 : ▲ CD147+5 CD55+4 CD24+3 CD16+3 CD29+2 CD45+1 CD64+1 CD62L+1 CD39+1 ▼ None |
| | | | 7 | 1362 | 4107095 : ▲ CD9+5 CD147+4 CD81+2 Siglec-8+2 CD29+2 CD45+1 CD55+1 CD24+1 CD99+1 CD39+1 ▼ None |
| | | | 8 | 876 | 4108095 : ▲ CD147+6 CD29+4 CD11c+3 CD45+2 CD55+2 HLA-DR+1 CD49d+1 CD64+1 CD14+1 CD38+1 CD56+1 CD99+1 CD39+1 ▼ None |
| | | | 9 | 117 | 4109095 : ▲ CD147+6 CD29+5 CD99+5 CD38+4 CD45+3 CD8a+3 CD49d+2 CD3+2 CD27+2 CD81+1 CD52+1 CD55+1 CD62L+1 CD56+1 ▼ None |
| | | | 10 | 93 | 41010095 : ▲ CD9+5 CD147+5 CD55+3 CD24+3 CD16+3 CD81+2 CD29+2 CD39+2 CD45+1 CD64+1 Siglec-8+1 CD38+1 CD99+1 ▼ None |

*Red MEM labels denote that the cell hotspot is a region with ≥95% expansion and blue MEM labels denote that the cell hotspot is a region with ≥95% contraction after T-REX analysis when comparing  Day 0 to Day 4 (±3 days)
