## Supplemental Table 2b for "Unsupervised machine learning reveals key immune cell subsets in COVID-19, rhinovirus infection, and cancer therapy"

**Supplemental Table 2b - MEM Labels for Enriched Features in CD4+ T cell Hotspots from COVID-19 Patients in Dataset 2**

| patient_id | degree_change | direction_change | DBSCAN_cluster | total_counts | Marker Enrichment Modeling (MEM) Label |
|---|---|---|---|---|---|
| COV24 | 2.053 | 1.000 | 1 | 81 | 2401095 : ▲ CD147+10 CD99+9 CD29+8 CD38+6 CD49d+4 CD3+4 CD81+2 CD52+2 CD56+2 CD28+2 CD55+1 CD137+1 CD62L+1 CD5+1 CD4+1 ▼ None |
| | | | 2 | 1 | 2402095 : ▲ CD99+10 CD147+9 CD29+8 CD3+5 CD27+5 CD38+5 CD28+3 CD81+2 CD52+2 CD62L+2 CD56+2 CD5+2 CD45+1 CD43+1 CD45RB+1 CD7+1 CD4+1 ▼ None |
| COV26 | N/A | N/A | N/A | N/A | N/A |
| COV27 | N/A | N/A | N/A | N/A | N/A |
| COV29 | 1.464 | 1.000 | 1 | 22 | 2901095 : ▲ CD147+10 CD29+8 CD99+8 CD38+7 CD49d+5 CD3+5 CD39+5 CD45+4 CD81+4 CD62L+4 CD28+4 CD55+3 CD56+3 HLA-DR+2 CD52+2 CD25+2 CD137+2 CD43+1 CD161+1 CD27+1 CD8a+1 CD5+1 CD4+1 ▼ None |
| | | | 2 | 186 | 2902095 : ▲ CD147+8 CD29+7 CD99+7 CD27+5 CD38+5 CD45+4 CD3+4 CD49d+3 CD52+3 CD28+3 CD81+2 CD62L+2 CD56+2 CD5+2 HLA-DR+1 CD43+1 CD55+1 CD161+1 CD137+1 CD8a+1 CD4+1 ▼ None |
| COV32 | 23.022 | -0.045 | 1 | 282 | 320105 : ▲ CD3+6 CD27+5 CD45RA+5 CD45+4 CD147+4 CD52+3 CD55+3 CD99+3 CD43+2 CD45RB+2 CD81+2 CD29+2 CD62L+2 CD127+2 CD38+2 CD5+2 CD28+1 CD8a+1 CD4+1 ▼ None |
| | | | 2 | 506 | 320205 : ▲ CD57+6 CD29+6 CD99+6 CD43+5 CD3+5 CD147+5 CD45+4 CD45RA+4 CD81+3 CD52+3 CD49d+2 CD5+2 CD45RB+1 CD127+1 CD38+1 CD56+1 CD8a+1 CD4+1 ▼ None |
| | | | 3 | 45 | 320305 : ▲ CD45+5 CD29+5 CD27+5 CD147+5 CD3+4 CD52+4 CD99+4 CD127+3 CD43+2 CD81+2 CD62L+2 CD5+2 CD45RB+1 CD55+1 CD38+1 CD28+1 CD8a+1 CD7+1 CD4+1 ▼ None |
| | | | 5 | 73 | 320505 : ▲ CD29+7 CD147+5 CD99+5 CD45+4 CD43+4 CD3+4 CD81+3 CD52+3 CD127+3 CD49d+2 CD5+2 CD161+1 CD137+1 CD56+1 CD8a+1 CD4+1 ▼ None |
| | | | 7 | 42 | 320705 : ▲ CD147+10 CD99+6 CD39+6 CD3+5 CD29+5 CD45+4 CD52+3 CD25+3 CD27+3 CD28+3 CD62L+2 CD38+2 CD56+2 CD5+2 CD43+1 CD55+1 CD4+1 ▼ None |
| | | | 4 | 73 | 3204095 : ▲ CD99+9 CD57+8 CD29+8 CD43+6 CD147+6 CD5+5 CD45+4 CD3+4 CD81+4 CD52+4 CD49d+3 CD45RB+1 CD161+1 CD137+1 CD127+1 CD56+1 CD8a+1 CD4+1 ▼ None |
| | | | 6 | 292 | 3206095 : ▲ CD99+10 CD29+9 CD147+9 CD27+7 CD45+5 CD3+5 CD52+5 CD38+5 CD49d+4 CD28+4 CD43+2 CD81+2 CD62L+2 CD56+2 CD5+2 CD4+2 CD45RB+1 CD55+1 CD161+1 CD137+1 CD127+1 CD8a+1 CD7+1 ▼ None |
| | | | 8 | 40 | 3208095 : ▲ CD99+7 CD3+6 CD27+6 CD147+5 CD45+4 CD81+4 CD52+4 CD45RA+4 CD43+3 CD45RB+3 CD55+3 CD29+3 CD7+3 CD62L+2 CD38+2 CD28+2 CD5+2 CD4+2 CD49d+1 CD127+1 CD8a+1 ▼ None |
| | | | 9 | 286 | 3209095 : ▲ CD99+9 CD57+8 CD29+8 CD43+6 CD147+6 CD3+5 CD56+5 CD81+4 CD52+4 CD45RA+4 CD45+3 CD49d+3 CD5+2 CD45RB+1 CD137+1 CD38+1 CD8a+1 CD4+1 ▼ None |
| | | | 10 | 44 | 32010095 : ▲ CD99+9 CD57+8 CD29+8 CD147+7 CD43+5 CD45+4 CD3+4 CD81+4 CD52+4 CD49d+3 CD38+2 CD5+2 CD45RB+1 CD161+1 CD137+1 CD127+1 CD56+1 CD8a+1 CD4+1 ▼ None |
| | | | 11 | 108 | 32011095 : ▲ CD45RA+8 CD99+7 CD3+6 CD27+6 CD147+5 CD45+4 CD45RB+4 CD52+4 CD43+3 CD81+3 CD55+3 CD29+3 CD38+3 CD7+3 CD62L+2 CD28+2 CD5+2 CD127+1 CD56+1 CD8a+1 CD4+1 ▼ None |
| | | | 12 | 22 | 32012095 : ▲ CD99+9 CD57+8 CD29+7 CD43+6 CD45RA+6 CD3+5 CD147+5 CD81+4 CD52+4 CD45+3 CD49d+3 CD45RB+2 CD5+2 CD137+1 CD56+1 CD8a+1 CD4+1 ▼ None |
| COV34 | 1.093 | 0.472 | 1 | 41 | 3401095 : ▲ CD99+10 CD29+9 CD27+8 CD147+6 CD127+5 CD45+4 CD43+4 CD3+4 CD52+4 CD62L+4 CD5+4 CD81+3 CD55+2 CD28+2 CD7+2 CD4+2 CD45RB+1 CD137+1 CD56+1 CD8a+1 ▼ None |
| | | | 3 | 26 | 3403095 : ▲ CD29+9 CD99+9 CD27+7 CD3+6 CD147+6 CD45+4 CD52+4 CD62L+4 CD45RA+4 CD43+3 CD45RB+3 CD81+3 CD55+2 CD127+2 CD28+2 CD8a+2 CD5+2 CD7+2 CD25+1 CD161+1 CD137+1 CD56+1 CD4+1 ▼ None |
| | | | 2 | 18 | 340205 : ▲ CD99+7 CD27+6 CD147+5 CD45+4 CD3+4 CD52+4 CD62L+4 CD38+3 CD43+2 CD45RB+2 CD81+2 CD55+2 CD29+2 CD127+2 CD8a+2 CD5+2 CD56+1 CD28+1 CD7+1 CD4+1 ▼ None |
| COV35 | N/A | N/A | N/A | N/A | N/A |
| COV36 | N/A | N/A | N/A | N/A | N/A |
| COV37 | N/A | N/A | N/A | N/A | N/A |
| COV39 | 0.474 | 1.000 | 1 | 44 | 3901095 : ▲ CD29+10 CD147+10 CD99+10 CD45+6 CD49d+4 CD3+4 CD27+4 CD52+3 CD28+3 CD38+2 CD81+1 CD137+1 CD62L+1 CD56+1 CD5+1 CD4+1 ▼ None |
| | | | 2 | 11 | 3902095 : ▲ CD99+10 CD29+9 CD147+8 CD45+6 CD3+4 CD49d+3 CD52+3 CD38+3 CD27+2 CD28+2 CD81+1 CD137+1 CD56+1 CD5+1 CD4+1 ▼ None |
| COV40 | N/A | N/A | N/A | N/A | N/A |
| COV41 | N/A | N/A | N/A | N/A | N/A |

*Red MEM labels denote that the CD4+ T cell hotspot is a region with ≥95% expansion and blue MEM labels denote that the CD4+ T cell hotspot is a region with ≥95% contraction after T-REX analysis when comparing Day 0 to Day 4 (±3 days)
