## Supplemental Table 3 for "Unsupervised machine learning reveals key immune cell subsets in COVID-19, rhinovirus infection, and cancer therapy"

**Supplemental Table 3: Fluorescent anitbody panel and tetramer selection for the analysis of RV-specific CD4+ T cells**

| Surface Antibody Panel | | | |
|---|---|---|---|
| **Fluorochrome** | **Marker** | **Clone** | **Vendor** |
| BB515 | CD45RO | UCHL1 | BD |
| FITC | Dump (CD14/CD19/CD8a | M5E2/HIB19/SK1 | Biolegend |
| Spark Blue 550 | CD3 | SK7 | Biolegend |
| PE | Tetramer 1 | NA | NA |
| PE-CF594 | Tetramer 2 | NA | NA |
| PE-Cy5 | Tetramer 3 | NA | NA |
| PerCP-Cy5.5 | CD95 | DX2 | Biolegend |
| PE-AF700 | CD25 | CD25-3G10 | Thermo Fisher |
| PE-Cy7 | CXCR5 | J252D4 | Biolegend |
| APC | Ki-67 | 20Raj1 | Thermo Fisher |
| AF647 | Tcf-1 | C63D9 | Cell Signaling Technologies |
| APC-Cy5.5 | CD38 | HIT2 | Thermo Fisher |
| Zombie Near IR | Live/Dead | NA | Biolegend |
| APC-Fire 750 | CCR7 | G043H7 | Biolegend |
| BUV 737 | CD45RA | H100 | BD |
| BV421 | CCR6 | G034E3 | Biolegend |
| Pacific Blue | ICOS | C398.4A | Biolegend |
| BD Horizon BV480 | CXCR3 | 1C6 | BD |
| BV570 | CD27 | O323 | Biolegend |
| BV605 | CCR5 | 2D7 | BD |
| BV650 | CD4 | OKT4 | Biolegend |
| BV711 | Tbet | 4B10 | Biolegend |
| BV750 | CD127 | HIL-7R-M21 | BD |
| BV786 | PD-1 | EH12.1 | BD |

| Tetramer Selection | | | | |
|---|---|---|---|---|
| **Subject ID** | **# Tets** | **PE Tetramer (DRB1)** | **PE-CF594 Tet (DRB1)** | **PE-Cy5 Tetramer (DRB1)** |
| RV001 | 3 | RV $VP2_{P3}$ (*0404) | RV $VP2_{P25}$ (*1501) | RV $VP1_{P18}$ (*0404) |
| RV002 | 3 | RV $VP2_{P24}$ (*0101) | RV $VP2_{P21}$ (*0301) | RV $VP1_{P23}$ (*0101) |
| RV003 | 1 | Empty | RV $VP2_{P25}$ (*1501) | Empty |
| RV004 | 2 | Empty | RV $VP1_{P18}$ (*0404) | RV $VP2_{P3}$ (*0404) |
| RV005 | 1 | Empty | RV $VP2_{P2}$ (*1101) | Empty |
| RV006 | 2 | Empty | RV $VP1_{P23}$ (*0701) | RV $VP2_{P26}$ (*0701) |
| RV007 | 2 | Flu $HA_{307-319}$ (*1101) | RV $VP2_{P2}$ (*1101) | Empty |
| RV008 | 1 | Empty | RV $VP2_{P21}$ (*0301) | Empty |

*For full details of rhinovirus tetramers refer to Muehling et al, JI 2016
